# Functional analysis of cyclic diguanylate-modulating proteins in *Vibrio fischeri*

**DOI:** 10.1101/2023.07.24.550417

**Authors:** Ruth Y. Isenberg, Chandler S. Holschbach, Jing Gao, Mark J. Mandel

## Abstract

As bacterial symbionts transition from a motile free-living state to a sessile biofilm state, they must coordinate behavior changes suitable to each lifestyle. Cyclic diguanylate (c-di-GMP) is an intracellular signaling molecule that can regulate this transition, and it is synthesized by diguanylate cyclase (DGC) enzymes and degraded by phosphodiesterase (PDE) enzymes. Generally, c-di-GMP inhibits motility and promotes biofilm formation. While c-di-GMP and the enzymes that contribute to its metabolism have been well-studied in pathogens, considerably less focus has been placed on c-di-GMP regulation in beneficial symbionts. *Vibrio fischeri* is the sole beneficial symbiont of the Hawaiian bobtail squid (*Euprymna scolopes*) light organ, and the bacterium requires both motility and biofilm formation to efficiently colonize. C-di-GMP regulates swimming motility and cellulose exopolysaccharide production in *V. fischeri*. The genome encodes 50 DGCs and PDEs, and while a few of these proteins have been characterized, the majority have not undergone comprehensive characterization. In this study, we use protein overexpression to systematically characterize the functional potential of all 50 *V. fischeri* proteins. All 28 predicted DGCs and 14 predicted PDEs displayed at least one phenotype consistent with their predicted function, and a majority of each displayed multiple phenotypes. Finally, active site mutant analysis of proteins with the potential for both DGC and PDE activities revealed potential activities for these proteins. This work presents a systems-level functional analysis of a family of signaling proteins in a tractable animal symbiont and will inform future efforts to characterize the roles of individual proteins during lifestyle transitions.

**IMPORTANCE:** C-di-GMP is a critical second messenger that mediates bacterial behaviors, and *V. fischeri* colonization of its Hawaiian bobtail squid host presents a tractable model in which to interrogate the role of c-di-GMP during animal colonization. This work provides systems-level characterization of the 50 proteins predicted to modulate c-di-GMP levels. By combining multiple assays, we generated a rich understanding of which proteins have the capacity to influence c-di-GMP levels and behaviors. Our functional approach yielded insights into how proteins with domains to both synthesize and degrade c-di-GMP may impact bacterial behaviors. Finally, we integrated published data to provide a broader picture of each of the 50 proteins analyzed. This study will inform future work to define specific pathways by which c-di-GMP regulates symbiotic behaviors and transitions.

## INTRODUCTION

Many bacteria exist in the environment in a free-living state, and upon encountering an animal host undergo dramatic developmental transitions. These adjustments enable symbiotic microbes–including mutualists, commensals, and pathogens–to acclimate to the physical, chemical, and nutritional milieu in the host; to resist immune responses; and to engage in behaviors required for survival and growth within the distinct host environment (1–5). To manage such transitions successfully, bacteria often inversely regulate motility and adhesion (6–11). In the motile state, it would be counterproductive to be adherent, and environmental bacteria use motility and chemotaxis to colonize novel niches, seek nutrition, and avoid predation (12–14). In contrast, adherent bacteria, especially those that have formed a multicellular biofilm, do not have a need for swimming motility. As a result there are multiple mechanisms that bacteria use to coordinately and inversely regulate these two broad behaviors (15–18). Alteration of the levels of the intracellular second messenger cyclic diguanylate (c-di-GMP) is a common mechanism used by bacteria to accomplish this purpose (19). In general, c-di-GMP promotes biofilm formation and inhibits motility (20). Enzymes that regulate c-di-GMP levels are diguanylate cyclases (DGCs), which synthesize c-di-GMP, and phosphodiesterases (PDEs), which degrade the molecule. DGCs contain GGDEF domains with conserved GG(D/E)EF active site residues, while PDEs contain EAL domains with conserved ExLxR active site residues or HD-GYP domains with conserved HD and GYP active site residues (21–26). Although many proteins have both GGDEF and EAL domains, only one domain is usually active, even when the amino acid motif for the other domain should function based on sequence conservation (27–30). However, environmental conditions may influence whether some dual-function proteins exhibit primarily DGC or PDE activity (31).

Much of what is known about c-di-GMP regulation of biofilm, motility, and host colonization is from studies on pathogenic species (32–34). Activation of cellulose production was the first role defined for c-di-GMP, in *Acetobacter xylinum* (now *Komagataeibacter xylinus*), and has remained one of most well characterized c-di-GMP-regulated phenotypes across taxa (20, 21). C-di-GMP-mediated cellulose production has been implicated in host colonization defects by pathogens *Escherichia coli* and *Salmonella enterica* serovar Typhimurium (33). Although much focus has been placed on defining roles for c-di-GMP in pathogenic associations, recent studies have focused on the impacts of c-di-GMP in bacteria during beneficial host associations. Host-derived ligands inactivate *Aeromonas veronii* DGC SpeD, which promotes host colonization (35). Additionally, high levels of c-di-GMP negatively impact the establishment of the symbiosis between the bioluminescent marine bacterium *Vibrio fischeri* and its host the Hawaiian bobtail squid (*Euprymna scolopes*) (36). *V. fischeri* is the sole, beneficial light organ symbiont of the Hawaiian bobtail squid (*Euprymna scolopes*), and the bacterium requires both swimming motility and biofilm formation to successfully colonize the host light organ (37–40). *V. fischeri* express polar flagella in seawater and form biofilm aggregates in the host mucus before migrating into the light organ (37, 38, 41–44). *V. fischeri* produce cellulose via the *bcs* locus-encoded cellulose synthase enzyme, which is activated by c-di-GMP, and genetic manipulation of c-di-GMP levels through deletion or overexpression of DGCs and PDEs modulates cellulose production (39, 45–48). Although cellulose is not required for symbiotic biofilm or squid colonization, an *in vivo* regulatory interaction exists between cellulose and the symbiosis polysaccharide (Syp) (36, 38, 39, 42, 49, 50).

We previously examined how global c-di-GMP levels impact bacterial behaviors in *V. fischeri*, including cellulose production, cellulose synthase and *syp* transcriptional reporter activities, and flagellar motility (36). For that work, we took advantage of a strain lacking seven DGCs and another strain lacking six PDEs to adjust the global c-di-GMP pool. That effort sparked our interest in the potential redundancy of the dozens of predicted c-di-GMP modulating enzymes and the capacity of the encoded proteins to impact bacterial behavior. It is common for bacteria, especially those with diverse lifestyles, to encode many DGCs and PDEs: *E. coli* encodes 29 such enzymes and *Vibrio cholerae* encodes 62 (51, 52). *V. fischeri* strain ES114 encodes 50 genes predicted to modulate c-di-GMP levels (**FIG. 1**) (53, 54). Of these possible c-di-GMP-modulating proteins, only a few have been characterized in depth for their roles in biofilm formation and/or swimming motility. DGCs MifA and MifB regulate magnesium-dependent motility by contributing to flagellar biogenesis (45). CasA is a DGC that is activated by calcium to inhibit motility and promote cellulose biofilm formation (48). Reduction of c-di-GMP levels by PDE BinA reduces cellulose synthesis (46). LapD has degenerate GGDEF and EAL active sites, one or both of which may recognize c-di-GMP to prevent cleavage of the biofilm-promoting adhesin LapV (47). In this model, the PDE PdeV activates biofilm dispersal by decreasing the c-di-GMP pool that activates LapD (47). In *V. fischeri* strain KB2B1, the ortholog of DGC VF_1200 was shown to inhibit swimming motility (55). The other 44 *V. fischeri* predicted c-di-GMP-modulating proteins have not undergone comprehensive phenotypic characterization, although a recent study was published where motility of c-di-GMP-modulating gene mutants were assessed (54). Additionally, none of the 50 c-di-GMP-modulating proteins have been shown to impact host colonization individually, although putative PDE VF_A0879 was predicted to be required for host colonization in a transposon insertion sequencing study (56). In this report, we systematically dissected the capability of *V. fischeri* predicted c-di-GMP-modulating enzymes to impact various biofilm and motility phenotypes through an approach combining systems-level overexpression analysis and targeted active site mutations.

**FIG 1.**
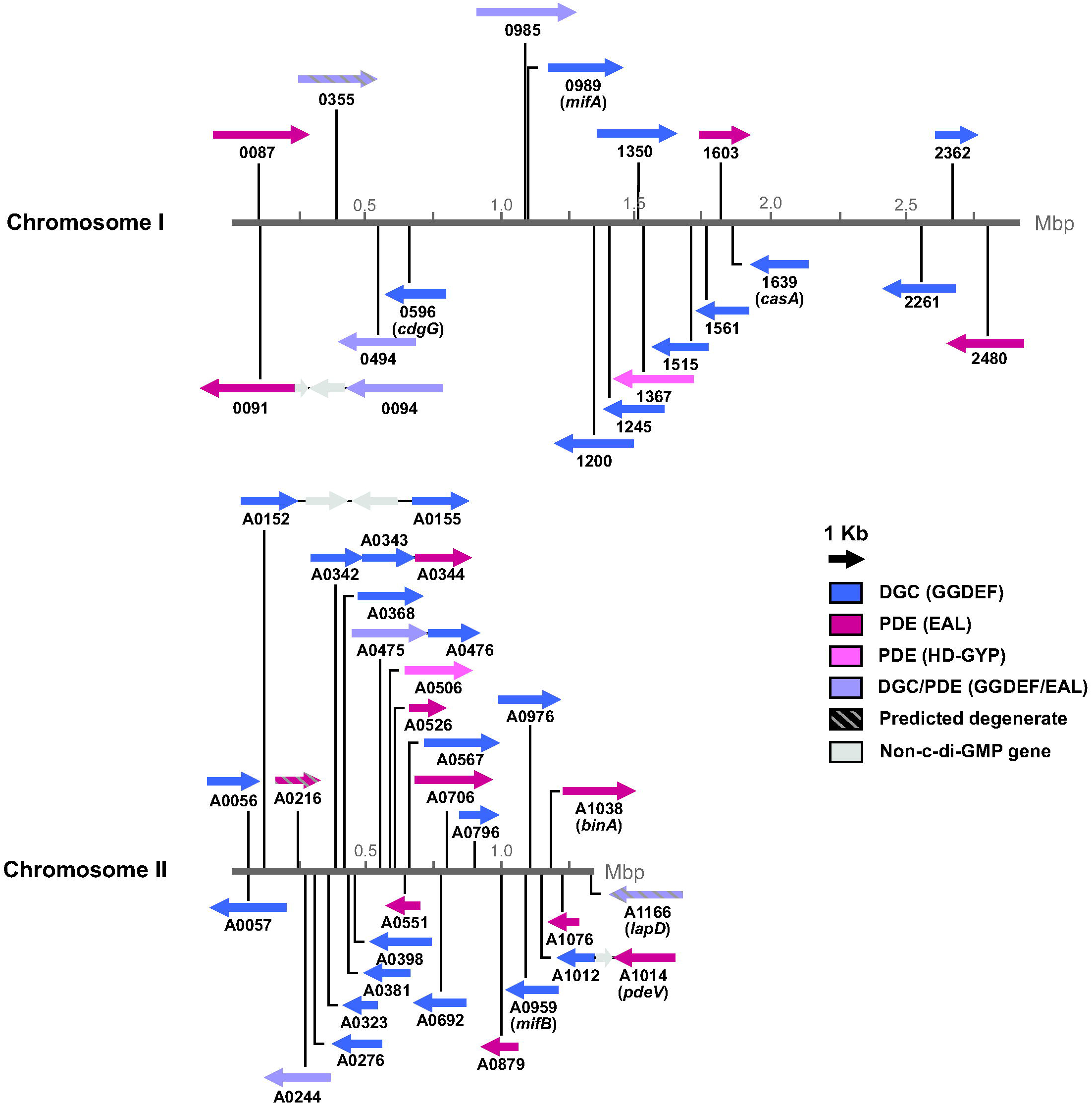
*V. fischeri* encodes 50 proteins across both chromosomes predicted to modulate c-di-GMP levels. The circular chromosomes are represented in a linear fashion for this representation. Numbers represent VF_ locus tags (e.g., VF_0087, VF_A0056, etc.).

## RESULTS

### *V. fischeri* encodes 50 proteins predicted to modulate c-di-GMP levels

The *V. fischeri* genome (strain ES114) encodes 50 proteins containing DGC and/or PDE domains (49, 50, 53, 54). Twenty-eight are predicted DGCs with GGDEF domains, 14 are predicted PDEs (12 with EAL domains, 2 with HD-GYP domains), 5 are predicted dual-function proteins with both GGDEF and EAL domains, and 3 are predicted to be nonfunctional due to degenerate active sites. The genes encoding these proteins are spread across both *V. fischeri* chromosomes, with most (31 out of 50) of the genes located on the smaller second chromosome (**FIG. 1**). To further characterize the functions of all 50 proteins, we took an overexpression approach to examine the function of each protein when individually overexpressed in *V. fischeri*. We sought this approach to be resilient against the redundancy we expect to exist within the large gene families. A similar overexpression approach has been effective to evaluate phenotypes of DGCs and PDEs in *V. cholerae* (57–61) and in other bacteria (28, 62–64). We used an IPTG-inducible vector to overexpress each protein in *V. fischeri* and performed assays to quantify cellulose production, swimming motility, and c-di-GMP levels. We also included control strains overexpressing VC1086 (a *V. cholerae* PDE) and QrgB (a *Vibrio campbellii* BB120 DGC), which have served as effective controls in multiple organisms (59, 61, 64–69). We note that the vector backbone used is the same as in many of the *V. cholerae* studies. Assays were conducted in a 96-well format to facilitate simultaneous, reproducible assays of the complete set of 54 test and control strains.

### Predicted *V. fischeri* DGCs and PDEs impact cellulose polysaccharide production

C-di-GMP promotes cellulose synthesis in many bacteria including *V. fischeri* (46, 48). To assess cellulose production across the set of proteins, we performed Congo red binding assays (46) of strains overexpressing each protein. Most DGCs (17/28 *V. fischeri* DGCs) increased cellulose production, consistent with their predicted function, including characterized DGCs MifA and CasA (**FIG. 2A**). Known *V. campbellii* DGC QrgB also increased cellulose production (**FIG. 2A**). The increase in cellulose production upon overexpression of known DGC MifA is consistent with published results (45), and the CasA overexpression result is consistent with data showing decreased cellulose production in a Δ*casA* strain (48). While overexpression of the remaining predicted DGCs did not significantly alter Congo red binding levels, many had a trend in the positive direction as expected and consistent with an increase in c-di-GMP levels (**FIG. 2A**).

**FIG 2.**
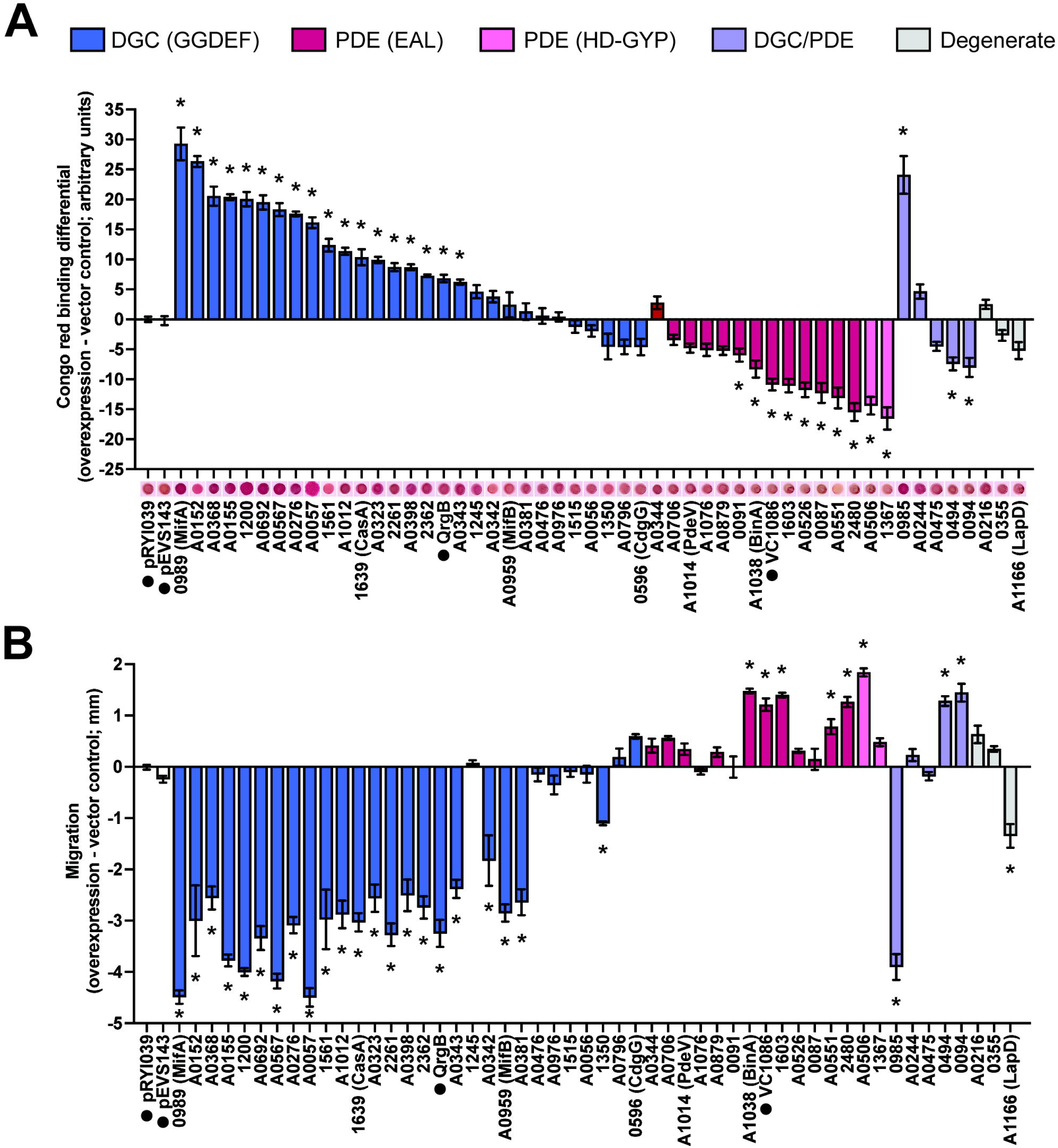
Many predicted *V. fischeri* DGCs and PDEs impact biofilm formation and swimming motility when overexpressed. **A.** Quantification of Congo red binding for *V. fischeri* strains overexpressing the indicated proteins relative to the pRYI039 empty vector control. For each strain, n = 3-8 biological replicates (24 for controls). Congo red images are representative. **B.** Quantification of migration through soft (0.3%) agar for *V. fischeri* strains overexpressing the indicated proteins relative to the pRYI039 empty vector control. For each strain, n = 4-11 biological replicates (33 for controls). For panels A and B, one-way analysis of variance (ANOVA) was used for statistical analysis, each bar represents the means of biological replicates, error bars represent standard errors of the mean, asterisks represent significance relative to the pRYI039 empty vector control (*, *P* < 0.05), numbers represent VF_ locus tags (e.g., VF_0087, VF_A0056, etc.); negative controls pRYI039 and pEVS143 as well as non-*V. fischeri* controls QrgB and VC1086 are also listed and indicated with a black dot.

Overexpression of 9/14 predicted PDEs significantly decreased cellulose production upon overexpression, consistent with their predicted function, including characterized PDE BinA. Known *V. cholerae* PDE VC1086 also decreased cellulose production (**FIG. 2A**). Our results are consistent with published results showing effects on cellulose production for PDE BinA (46, 47).

*V. fischeri* encodes five proteins with both GGDEF and EAL domains. Proteins that contain both GGDEF and EAL domains typically only have one domain exhibit activity, even if the amino acid motif for the other domain is conserved (27–30), though there is evidence of a dual-function protein capable of exhibiting either DGC or PDE activity depending on conditions (31). We therefore expected these dual-function proteins to behave predominantly as DGCs or PDEs. Of the five predicted dual-function proteins, overexpression of VF_0985 increased cellulose production while VF_0094 and VF_0494 decreased cellulose production compared to the empty vector control (**FIG. 2A**).

Finally, we examined the three predicted “degenerate” c-di-GMP modulating enzymes; i.e., proteins with intact GGDEF and/or EAL domains but with degenerate GG(D/E)EF and/or ExLxR active site motifs that are not predicted to function. None of these proteins significantly impacted cellulose production (**FIG. 2A**).

### Predicted *V. fischeri* DGCs and PDEs influence flagellar motility

As a second behavioral output of c-di-GMP levels, we proceeded to assay swimming motility in the same set of strains. We conducted swimming motility assays of strains overexpressing each protein in TBS soft agar and in TBS soft agar with the addition of either magnesium or calcium, which are known to influence swimming behavior by *V. fischeri* (54, 70). We observed that most DGCs (21/28) and 6/14 PDEs showed the expected overexpression results of inhibiting or promoting motility, respectively, including known *V. fischeri* DGCs MifA, MifB, and CasA and known PDE BinA (**FIG. 2B**). *V. campbellii* DGC QrgB and *V. cholerae* PDE VC1086 also showed the expected results (**FIG. 2B**). Overexpression of either DGC MifA, MifB, or CasA diminished motility under all conditions tested despite having known cation-specific motility phenotypes (**FIG. 2B; FIG. S1**). MifA and MifB inhibit motility in the presence of magnesium (45), while CasA inhibits motility in the presence of calcium (48), and we suspect that overexpression of the proteins likely amplified their respective DGC activities and bypassed the requirement for the respective cations. One example where we observed a media-specific effect is for VF_0985, which strongly inhibited motility upon overexpression in TBS and TBS-Ca, relative to the empty vector control, but not in TBS-Mg (**FIG. 2B; FIG. S1**).

Overexpression of predicted degenerate proteins VF_0355 and VF_A0216 did not significantly impact motility (**FIG. 2B; FIG. S1**). Overexpression of LapD decreased motility in two of the three motility media types (**FIG. 2B; FIG. S1**), but this is likely independent of c-di-GMP modulation and is consistent with the function of LapD in inhibiting biofilm dispersal (47), which is associated with decreased motility.

### Expression level of *V. fischeri* proteins does not correlate with magnitude of phenotypes

Of the *V. fischeri* proteins assayed, 13 did not have significant phenotypes for either cellulose production or motility: DGCs CdgG, VF_1245, VF_1515, VF_A0056, VF_A0476, VF_A0796; PDEs VF_A0706, VF_A0879, VF_A1076, and PdeV; dual-function proteins VF_A0244 and VF_A0475; and predicted degenerate proteins VF_0355 and VF_A0216 (**FIG. 2; FIG. S1**). This result for these proteins could be due to lack of enzymatic activity under the conditions tested, or could be the result of posttranscriptional regulation that prevents these proteins from being expressed. To test whether the lack of significant phenotypes from these proteins was due to protein expression, we assessed expression of several FLAG-tagged proteins via western blot. We selected proteins with strong phenotypes in both the cellulose and motility assays (MifA, VF_A0057, and VF_A0155), one protein with a phenotype in just the motility assay (VF_A0342), proteins with no significant phenotypes in either assay (VF_1515, VF_A0056, and VF_A0476), and a predicted degenerate protein with no significant phenotypes (VF_A0216). All three proteins with strong phenotypes in both assays were expressed robustly (**FIG. 3**). Two of the proteins with no significant phenotypes (VF_A0056 and VF_A0476) were expressed, with the intensity of the band for VF_A0056 as strong as the bands for the proteins with significant phenotypes in both assays (**FIG. 3**). The predicted degenerate protein VF_A0216 was also expressed, as was VF_A0342 which only had a significant motility phenotype (**FIG. 3**).

**FIG 3.**
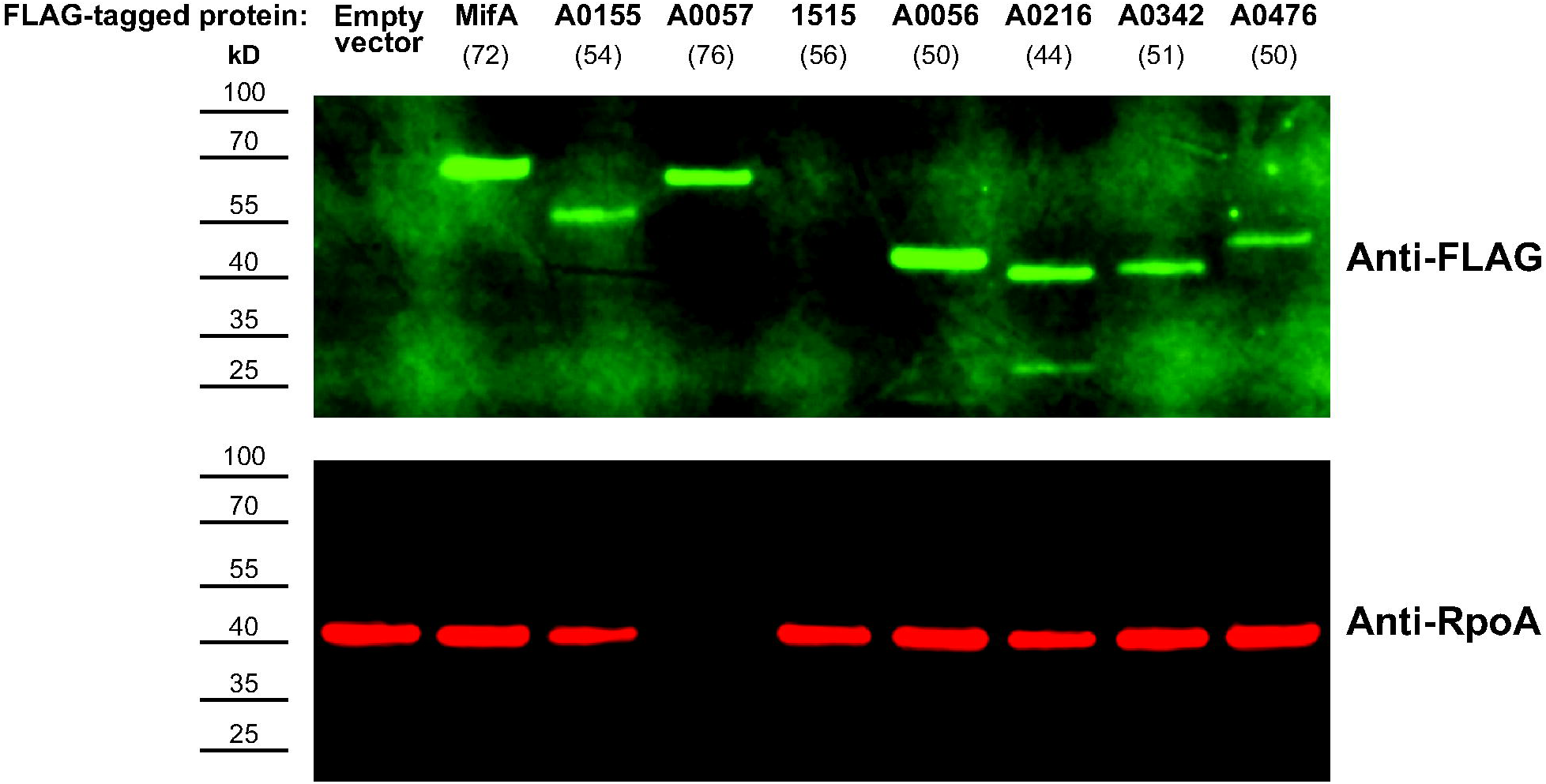
Most *V. fischeri* proteins tested are expressed from the overexpression vector. Western blot of whole-cell lysates of *V. fischeri* expressing indicated FLAG-tagged proteins from the pEVS143 vector. Predicted band sizes (kD) for each protein are indicated in parentheses. Anti-FLAG Rabbit IgG was used as the primary antibody and LI-COR IRDye 800CW Goat anti-Rabbit IgG was used as the secondary antibody. Anti-RpoA Mouse IgG was used as a loading control, binding the RNAP α subunit, and LI-COR IRDye 680RD Goat anti-mouse IgG was used as the secondary antibody of the loading control. Western blot is representative of n = 3 biological replicates.

VF_1515, which had no significant phenotypes, was the only protein tested that had no observable expression via western blot, suggesting that VF_1515 protein is not produced at a substantial level under the conditions tested (**FIG. 3**). Therefore, while we expect that most proteins expressed from pEVS143 are found at appreciable levels in the cell, including those proteins that do not yield detectable phenotypes, in at least one case we found that the protein assayed was not expressed stably in the cell. Further, the magnitude of the cellulose and motility phenotypes do not correlate with protein expression level for proteins that are expressed.

### Predicted *V. fischeri* DGCs and PDEs modulate c-di-GMP reporter levels

Congo red and motility assays revealed that most of the *V. fischeri* DGCs and PDEs influence phenotypes associated with c-di-GMP. We next asked to what capacity these proteins are capable of modulating c-di-GMP levels using the pFY4535 fluorescent c-di-GMP reporter plasmid (71), which we used previously to quantify c-di-GMP levels in *V. fischeri* (36). We expected overexpression of DGCs to increase c-di-GMP levels and overexpression of PDEs to decrease c-di-GMP levels relative to the empty vector control. We also expected proteins with the strongest cellulose and/or motility phenotypes to have the strongest effects of c-di-GMP reporter activity. Consistent with their predicted function, overexpression of all 28 predicted DGCs increased c-di-GMP reporter activity, including characterized *V. fischeri* DGCs MifA and MifB (**FIG. S2A**). The results for overexpression of MifA and MifB are consistent with published results (45). In contrast to the results with DGCs, upon overexpression only 4/14 predicted PDEs, VF_0087, VF_A0506, VF_A0526, and VF_A0706, had the expected effect of significantly decreasing c-di-GMP reporter activity (**FIG. S2A**). These results are also in contrast to the cellulose and motility results, where overexpression of the majority of PDEs elicited at least one of the expected phenotypes, and no PDEs had significant opposite cellulose or motility phenotypes (**FIG. 2; FIG. S2D**). BinA is a characterized PDE that negatively influences c-di-GMP levels (absence of BinA resulted in increased c-di-GMP compared to WT) (46). Our results show that BinA overexpression diminished c-di-GMP reporter activity by 57%, but this result was not statistically significant (**FIG. S2A**). However, the cellulose and motility results for BinA are among the strongest of the PDEs (**FIG. 2; FIG. S2D**). Similarly, overexpression of known *V. cholerae* PDE VC1086 also did not significantly alter c-di-GMP reporter activity (**FIG. S2A**), but did significantly alter cellulose production and motility in the expected directions (**FIG. 2; FIG. S2D**). Surprisingly, five predicted PDEs, VF_0091, VF_1367, VF_A0344, and VF_A0879, and VF_A1076 significantly increased c-di-GMP reporter activity upon overexpression (**FIG S2A**). The same results were observed when the strains were assayed individually rather than in the arrayed 96-well format, with the exception of VF_1367 which did not have significant results in the individual assay format (**FIG. S2B**). Of these five PDEs, VF_0091 significantly decreased cellulose production and none significantly increased motility, but these phenotypes do not correspond with increased c-di-GMP reporter activity (**FIG. 2; FIG. S2D**). One interpretation of these results is that the c-di-GMP reporter reports a local level of c-di-GMP while the Congo red and motility assays report global levels. Local c-di-GMP levels measured by the reporter may correspond with global levels for the majority of strains, but not for strains overexpressing some PDEs, where the dynamic range of c-di-GMP levels is much more limited and local pools may be subject to substantial variation from the global dynamics. We previously reported that in a high c-di-GMP strain of *V. fischeri*, overexpression of PDE VC1086 reduced c-di-GMP reporter activity 1.6-fold when measured with the same reporter (36). However, overexpression with the same vector in wild-type *V. fischeri* had no effect on the already-low c-di-GMP reporter activity. Therefore, while the c-di-GMP reporter levels were well-correlated with c-di-GMP-responsive phenotypes for the DGCs, data from the reporter were not informative for predicting phenotypes from the low basal wild-type levels. To test whether the pFY4535 c-di-GMP biosensor could better report decreased c-di-GMP in a high c-di-GMP background, we overexpressed select PDEs in a strain of *V. fischeri* deleted for six PDEs (Δ6PDE) and has been demonstrated to have high basal c-di-GMP levels (36). Every PDE tested diminished c-di-GMP reporter activity in the high c-di-GMP background, including VF_0091 and VF_A1076, which exhibited the highest reporter activity in the wild-type background (**FIG. S2C**). Dual-function protein VF_0094 diminished reporter activity comparable to the strongest PDE in the assay (**FIG. S2C**), consistent with the cellulose and motility results for this protein in the wild-type background (**FIG. 2**). To further probe the accuracy of the c-di-GMP reporter for functional characterization of DGC and PDE activities, we selected several PDE overexpression strains to measure c-di-GMP levels using a commercially available ELISA kit. The results of the ELISA c-di-GMP quantification did not agree with the phenotypic characterization of the selected PDEs, nor did they agree with the c-di-GMP reporter data (**FIG. S3**). We also quantified the phenotypic correlation of the reporter data with our Congo red (R^2^ = 0.69) and motility (R^2^ = 0.66) results, which was lower in both cases than the correlation of the Congo red and motility phenotypes to each other (R^2^ = 0.77). Given these questions about the utility of the reporter activity in the wild-type background when c-di-GMP levels are diminished, for our overall question of the catalytic potential of the proteins assayed we argue that in this case it is most prudent to infer global c-di-GMP-dependent processes from the phenotypic assays of cellulose production and swimming motility. In the next section, we test this hypothesis through mutation of relevant c-di-GMP active sites.

### Select *V. fischeri* DGCs and PDEs impact motility and biofilm phenotypes through their c-di-GMP catalytic sites

Most of the DGCs and PDEs exhibited the expected cellulose and motility phenotypes upon overexpression (**FIG. 2**). If the results we observed were due to catalytic function, then mutation of the active sites should impact these phenotypes. Changing the second amino acid in the GGDEF motif to A (62, 72, 73) drastically reduced the effects of MifA, VF_A0057, VF_A0152, and VF_A0398 on cellulose production and motility (**FIG. 4**). These results confirm that intact GGDEF domains are required for the regulatory functions of these proteins.

**FIG 4.**
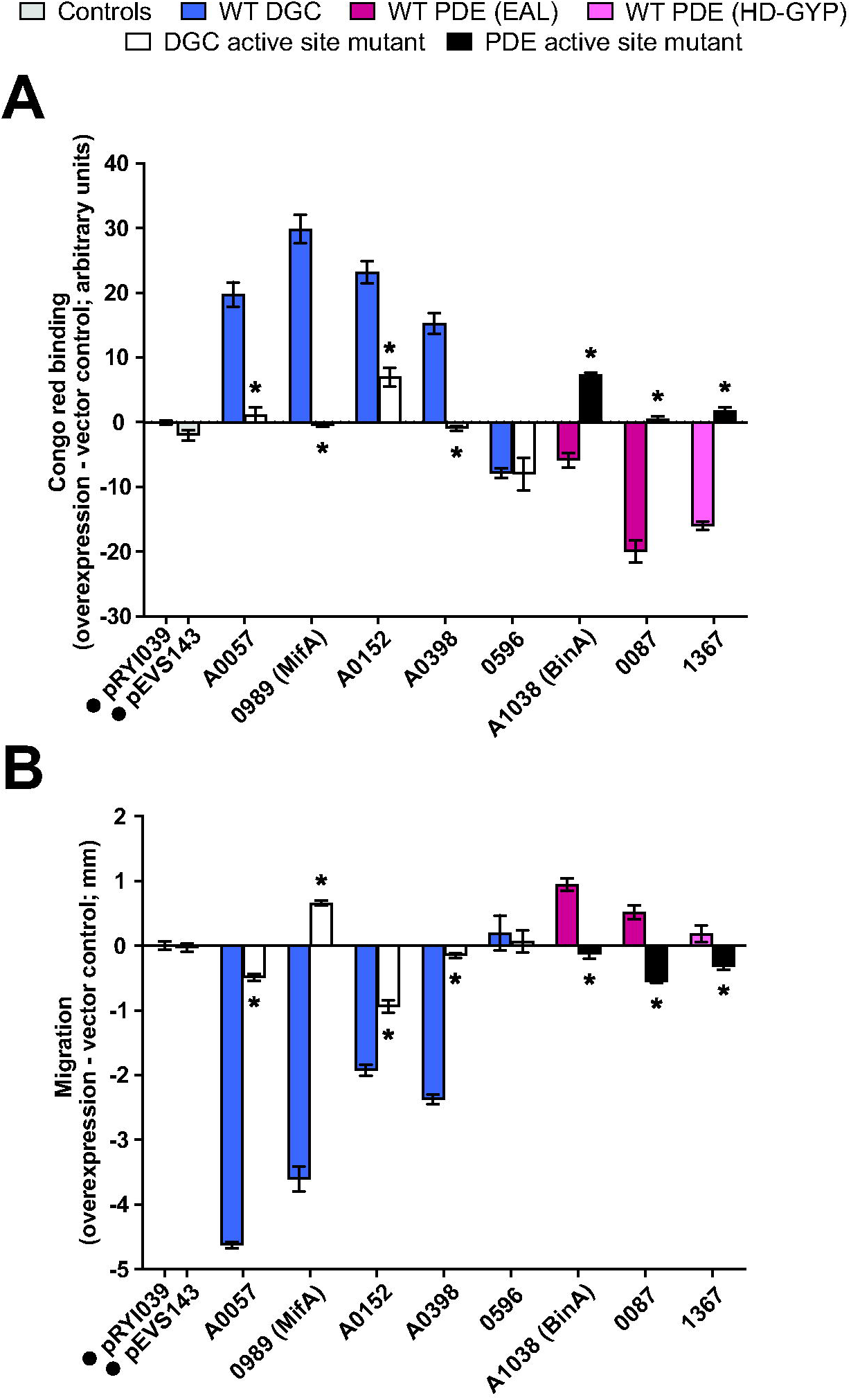
Active site residues are required to modulate cellulose production and motility for selected DGCs and PDEs. **A.** Quantification of Congo red binding for *V. fischeri* strains overexpressing the indicated proteins relative to the pRYI039 empty vector control. For each strain, n = 4-5 biological replicates (10 for controls). **B.** Quantification of migration through soft (0.3%) agar for V. fischeri strains overexpressing the indicated proteins relative to the pRYI039 empty vector control. For each strain, n = 5 biological replicates (10 for controls). For A and B, unpaired t tests were used for statistical analysis, each bar represents the means of biological replicates, error bars represent standard errors of the mean, asterisks represent significance of a mutant relative to the corresponding wildtype protein (*, *P* < 0.01) and numbers represent VF_ locus tags (e.g., VF_0087, VF_A0056, etc.); negative controls pRYI039 and pEVS143 are indicated with a black dot.

VF_A0057 is one of the more interesting predicted DGCs because it has a non-consensus GGDEF domain with a serine instead of a glycine in the first position of the motif (i.e., SGDEF). The altered motif is not functional in *V. cholerae* CdgG, which led us to predict it to be similarly nonfunctional in *V. fischeri* VF_A0057 (72). Surprisingly, though, mutation of the second amino acid of the motif in VF_A0057, resulting in the amino acid sequence SADEF, substantially mitigated the effects of the protein on cellulose production and motility, suggesting that the non-canonical SDGEF motif in this protein is functional and that VF_A0057 likely has DGC activity (**FIG. 4**). CdgG, a homolog of the nonfunctional *V. cholerae* protein CdgG, is another predicted DGC with a non-consensus active site where the first amino acid is a serine: SGEEF. However, overexpression of CdgG or the variant (SAEEF) did not promote cellulose production or inhibit motility (**FIG. 4**).

To examine the catalytic site of the PDEs, we similarly replaced the first residue of the ExLxR motif with an alanine, a mutation that has been used previously to disrupt EAL domain activity (46, 59, 74, 75). AxLxR mutants of BinA and VF_0087 attenuated the effects of these proteins on cellulose production and motility (**FIG. 4**). These results confirm previous work (46, 54, 55), and are the first demonstration of the requirement for the active site residues for VF_0087 function.

VF_1367 is one of two predicted HD-GYP domain PDEs encoded by *V. fischeri*. The motif encoded is HD-GYL, but the presumed HD catalytic dyad for HD superfamily proteins is still intact (23). Here, we changed the second amino acid to an alanine, which has been used to disrupt HD-GYP domain activity (25, 60). Overexpression of an HA-GYL variant of VF_1367 obviated the impact on cellulose production, which was the major phenotype observed on overexpression of the wild-type protein (**FIG. 4**). Overexpression of active site mutants of predicted and characterized DGCs and PDEs thus supports the predicted biochemical activities of individual proteins and validates our overall approach of overexpressing predicted c-di-GMP-modulating proteins to understand the genome-wide functional landscape.

### Challenges in assigning functions for dual-domain proteins

Three of the five *V. fischeri* dual-function proteins had clear phenotypes in both cellulose production and motility assays, suggesting they may primarily function as DGCs or PDEs (**FIG. 2**). However, overexpression of proteins with both GGDEF and EAL domains can only hint at whether a protein primarily has DGC or PDE activity and does not discern which domains are active/inactive. To assess whether one, both, or neither of the DGC and PDE domains contribute to the effects of these predicted dual-domain protein, we assayed strains overexpressing GGDEF mutants, EAL mutants, or GGDEF/EAL double mutants of each protein. As one example, phenotypic assays strongly suggested that VF_0985 behaved as a DGC given its impact on cellulose and motility (**FIG. 2**). A VF_0985 AAL mutant behaved like the wild-type protein in all assays when overexpressed, while a GADEF mutant drastically reduced the effects of VF_0985 on cellulose production and motility (**FIG. 5**). Interestingly, mutation of the GGDEF domain seemed to reverse the effects of VF_0985: the GGDEF mutant behaved similar to a PDE in both assays (**FIG. 5**), suggesting the EAL domain may be active. When the VF_0985 double GADEF/AAL mutant was overexpressed, cellulose production resembled the single GGDEF mutant, while little effect on motility was observed (**FIG. 5**). These results suggest that, under the conditions tested, VF_0985 likely functions predominantly as a DGC, but that we may be detecting cryptic PDE activity in the variant in which DGC activity is absent. Active site mutant analysis of the remaining four dual-function proteins similarly suggest a more complicated picture than one dominant domain. Further in-depth studies of these proteins will be required to dissect their roles in cellulose production and motility, and the mechanisms by which these proteins perform those roles.

**FIG 5.**
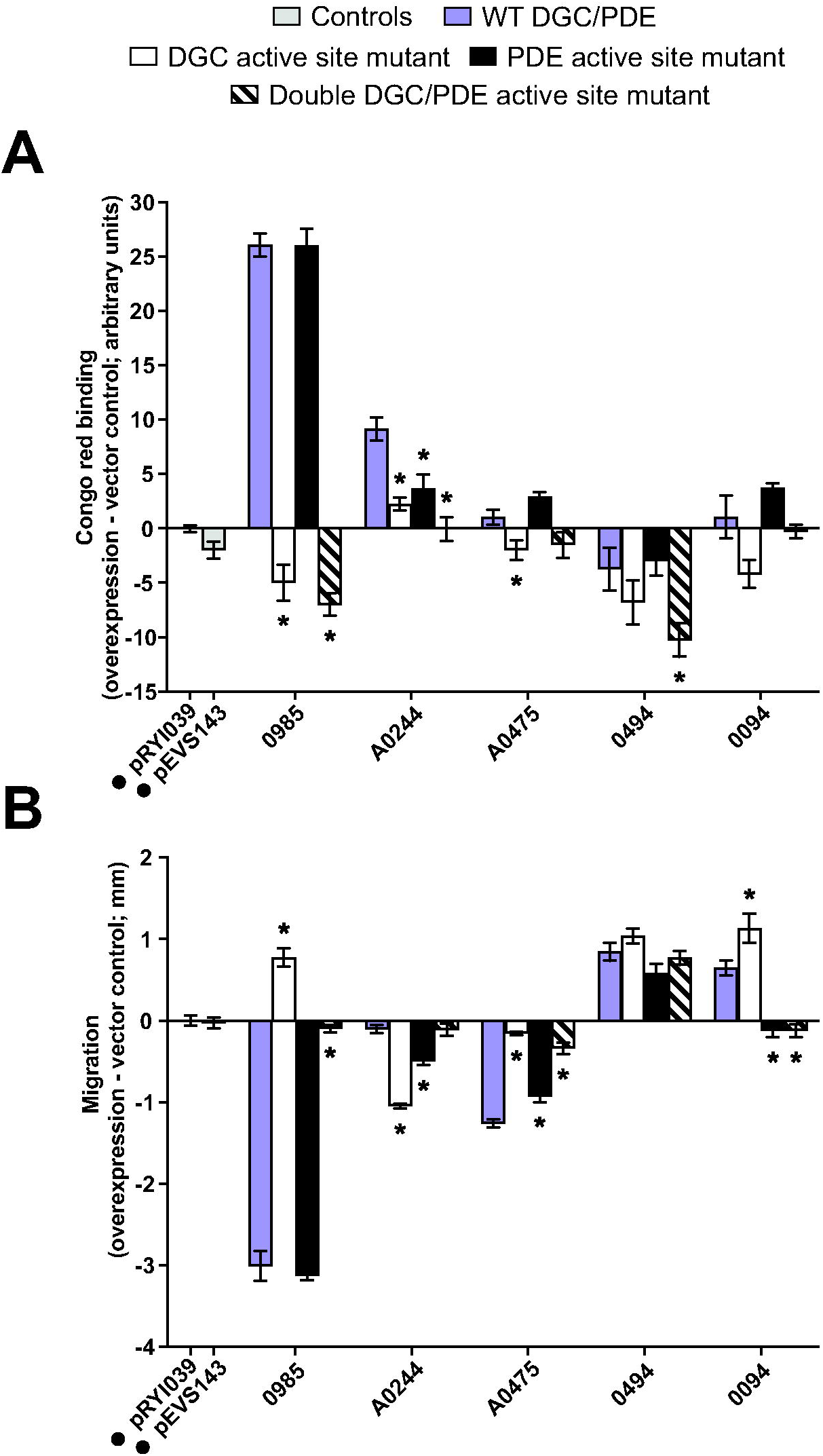
VF_0985 is the only dual-function protein with strong active site-dependent phenotypes. **A.** Quantification of Congo red binding for *V. fischeri* strains overexpressing the indicated proteins relative to the pRYI039 empty vector control. For each strain, n = 4-5 biological replicates (10 for controls). **B.** Quantification of migration through soft (0.3%) agar for *V. fischeri* strains overexpressing the indicated proteins relative to the pRYI039 empty vector control. For each strain, n = 5 biological replicates (10 for controls). For A and B, unpaired t tests were used for statistical analysis, each bar represents the means of biological replicates, error bars represent standard errors of the mean, asterisks represent significance of a mutant relative to the corresponding wildtype protein (*, *P* < 0.05) and numbers represent VF_ locus tags (e.g., VF_0087, VF_A0056, etc.); negative controls pRYI039 and pEVS143 are indicated with a black dot.

## DISCUSSION

*V. fischeri* is an emerging model for studies of c-di-GMP regulation, and especially for how c-di-GMP impacts animal host colonization. We previously demonstrated that high global levels of c-di-GMP impair colonization, though it is unknown how levels remain low to facilitate a productive host-microbe symbiosis. An ongoing goal is to elucidate relevant signaling that enables proper c-di-GMP in the host and during transitions to and from the host-associated state. Therefore, the goal of the current study was to identify which of the predicted 50 c-di-GMP modulating proteins–DGCs, PDEs, dual-function DGC/PDEs, and likely degenerate enzymes–are able to elicit c-di-GMP-responsive phenotypes in *V. fischeri*. In **Figure 6** we assembled the results from our analyses to demonstrate cellulose production and swimming motility results for all 50 *V. fischeri* proteins and the two control proteins. Furthermore, we integrated data from a recent study (54) that examined swimming motility of single-gene mutants in the same *V. fischeri* proteins. The resulting table (**FIG. 6**) presents a powerful visualization of current knowledge of the catalytic potential of these gene families. Conducted in different labs, there is remarkable consistency among the results. There are more proteins with significant effects observed upon overexpression (32/50 in at least one motility assay) compared to gene deletion (20/50), which was expected, and supports our approach to use the overexpression approach to reveal function in a family where redundancy is expected. The one case in which discrepancies were noted were for DGC VF_A0368, where we had consistent results across all conditions and the deletion approach yielded significant results only on TBS swim plates. The only cases in which a phenotype was observed solely in the deletion study were for DGCs CdgG and VF_A0476 as well as predicted degenerate protein VF_0355, where Δ*cdgG*, Δ*VF_A0476*, and Δ*VF_0355* strains each yielded the unexpected phenotype of decreased motility and we did not observe any significant cellulose or motility phenotypes upon overexpression of these factors.

**FIG 6.**
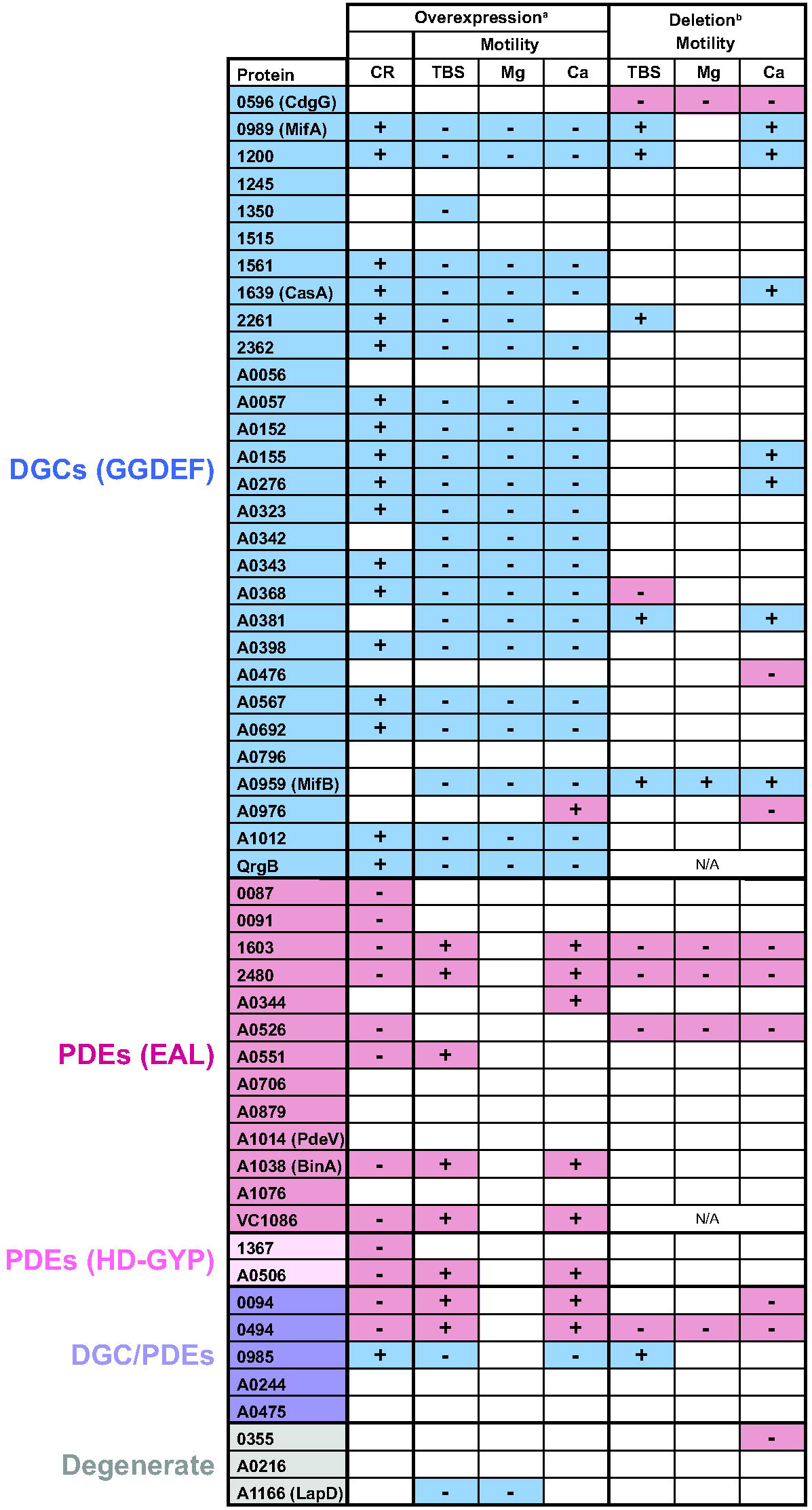
Integration of phenotypic data for the *V. fischeri* DGCs and PDEs. Numbers represent VF_ locus tags (e.g., VF_0087, VF_A0056, etc.); non-*V. fischeri* controls QrgB and VC1086 are also listed. Overexpression data are from this study; deletion data are integrated from Shrestha *et al*., 2022. Blue coloring indicates phenotypes expected from elevated c-di-GMP, whereas pink indicates phenotypes expected from reduced c-di-GMP. White indicates no significant change. ^a^This study, ^b^Shrestha *et al.* 2022.

The proteins we observed to exhibit the strongest phenotypes were DGCs VF_1200, VF_A0057, VF_A0152, VF_A0155, VF_A0276, VF_A0368, VF_A0567, and VF_A0692; PDEs VF_1603, VF_2480, VF_A0506, and VF_A0551; and dual-function protein VF_0985 (**FIG. 2 and 6**). We also identified interesting phenotypes in VF_0985, which inhibits motility only when magnesium is absent. For all of these proteins, little work has been conducted on a mechanistic basis, and our study presents candidates to pursue that may mediate relevant signaling in the host or during key lifestyle transitions. A benefit of the overexpression approach is that it can reveal phenotypes that are not apparent during single deletions. For example, a protein that is expressed during host colonization or in a specific environmental condition, but not during culture growth, may be assayed for function in this manner. Conversely, a limitation of our method is that the likely higher levels of each examined protein obscures natural regulation that will certainly be relevant to understand signaling *in vivo*. Therefore, this work pares down a complex family of 50 proteins to a narrower set for more focused individual studies.

While the data in **Figure 6** are remarkably consistent, it is clear that there are situations in which protein overexpression impacts motility and not cellulose, or motility in some media and not others. In fact, on a fine scale there are 11 different patterns to the overexpression data among *V. fischeri* proteins in **Figure 6**. This result is even more remarkable given that there was no difference in transcriptional regulation in our experimental setup. Although we detected distinct protein levels that likely suggest posttranscriptional regulation (**FIG. 3**), steady state protein levels did not correlate with the magnitude of the cellulose or motility phenotypes observed.

Therefore, our results suggest that local signaling effects—e.g., localization, protein-protein interactions—may play a major role in how these factors mediate host interactions. The factors that impacted motility in the expected direction (but did not have a significant affect on cellulose production) were DGCs VF_1350, VF_A0342, VF_A0381, and MifB. Conversely, cellulose production (but not motility) was impacted on overexpression of PDEs VF_0087, VF_0091, VF_1367, and VF_A0526. In the case of the DGCs, all four are predicted to have transmembrane domain(s) (54), raising the possibility that membrane localization, and perhaps localization at the flagellar pole may impact the motility-specific phenotype.

The effects of local signaling and protein-protein interactions by c-di-GMP-modulating proteins is particularly exemplified by dual-function proteins as has been demonstrated across Gram-negative bacteria. *Pseudomonas aeruginosa* RmcA encodes GGDEF and EAL domains, both of which are functional, but subcellular localization of RmcA mediated by interaction with the response regulator CbrB activates RmcA PDE activity, subsequently reducing type III secretion system gene expression and increasing biofilm formation (30). *V. cholerae* dual-function protein MbaA interacts with the periplasmic binding protein NspS, which senses norspermidine and activates MbaA DGC activity to increase the local c-di-GMP pool (76, 77). C-di-GMP synthesized by MbaA binds to specific high-affinity biofilm effectors when norspermidine levels are low, sensitizing the cell to norspermidine (77). *Pseudomonas fluorescens* encodes 43 proteins that metabolize c-di-GMP, and when tested across a broad range of conditions it was determined that ligand signaling, protein-protein interactions, and/or transcriptional regulation are all central to c-di-GMP signaling, thus highlighting the importance of local c-di-GMP signaling (78). This backdrop of growing evidence for local signaling and regulated activity of distinct DGCs and PDEs provide hints as to where such regulation could occur in *V. fischeri* given the unique patterns in our data. Recent publication of a more sensitive reporter that is amenable to single-cell imaging may be a valuable tool to investigate both the levels and subcellular dynamics of c-di-GMP in *V. fischeri* (79).

We previously demonstrated that global *V. fischeri* c-di-GMP levels need to be kept sufficiently low to successfully colonize squid (36), but the specific signals and pathways involved are yet to be determined. During colonization initiation, *V. fischeri* responds to squid-derived signals to help guide them to the light organ (37, 80–83). Therefore it is likely that one or more squid-specific signals may be sensed by c-di-GMP-modulating proteins to elicit effects on certain biofilm and/or motility pathways. Our study presents a set of DGCs and PDEs that are demonstrably functional in multiple assays in *V. fischeri*, and provide strong candidates as proteins to regulate c-di-GMP levels to facilitate functional and reproducible squid host colonization.

## MATERIALS AND METHODS

### Bacterial strains, plasmids, and media

*V. fischeri* and *E. coli* strains used in this study are listed in Table 1. Plasmids used in this study are listed in Table 2. *V. fischeri* strains were grown at 25°C in Luria-Bertani salt (LBS) medium (per liter: 25 g Difco LB broth [BD], 10 g NaCl, 50 mM Tris buffer [pH 7.5]) or tryptone broth salt (TBS) medium (per liter: 10 g Bacto Tryptone [Gibco], 20 g NaCl, 35 mM MgSO_4_ [where noted], 10 mM CaCl_2_ [where noted], 50 mM Tris buffer [pH 7.5]) where noted. *E. coli* strains used for cloning and conjugation were grown at 37°C in Luria-Bertani (LB) medium (per liter: 25 g Difco LB broth [BD]). When needed, antibiotics were added to the media at the following concentrations: kanamycin, 100 μg/mL for *V. fischeri* and 50 μg/mL for *E. coli*; gentamicin, 2.5 μg/mL for *V. fischeri* and 5 μg/mL for *E. coli*. When needed, 100 μM isopropyl ß-D-1-thiogalactopyranoside (IPTG) was added to the media. For Congo red media, 40 μg/mL Congo red and 15 μg/mL Coomassie blue were added to LBS. When needed, growth media was solidified using 1.5% agar. Plasmids were introduced from *E. coli* strains into *V. fischeri* strains using standard techniques (84, 85).

**Table 1.**
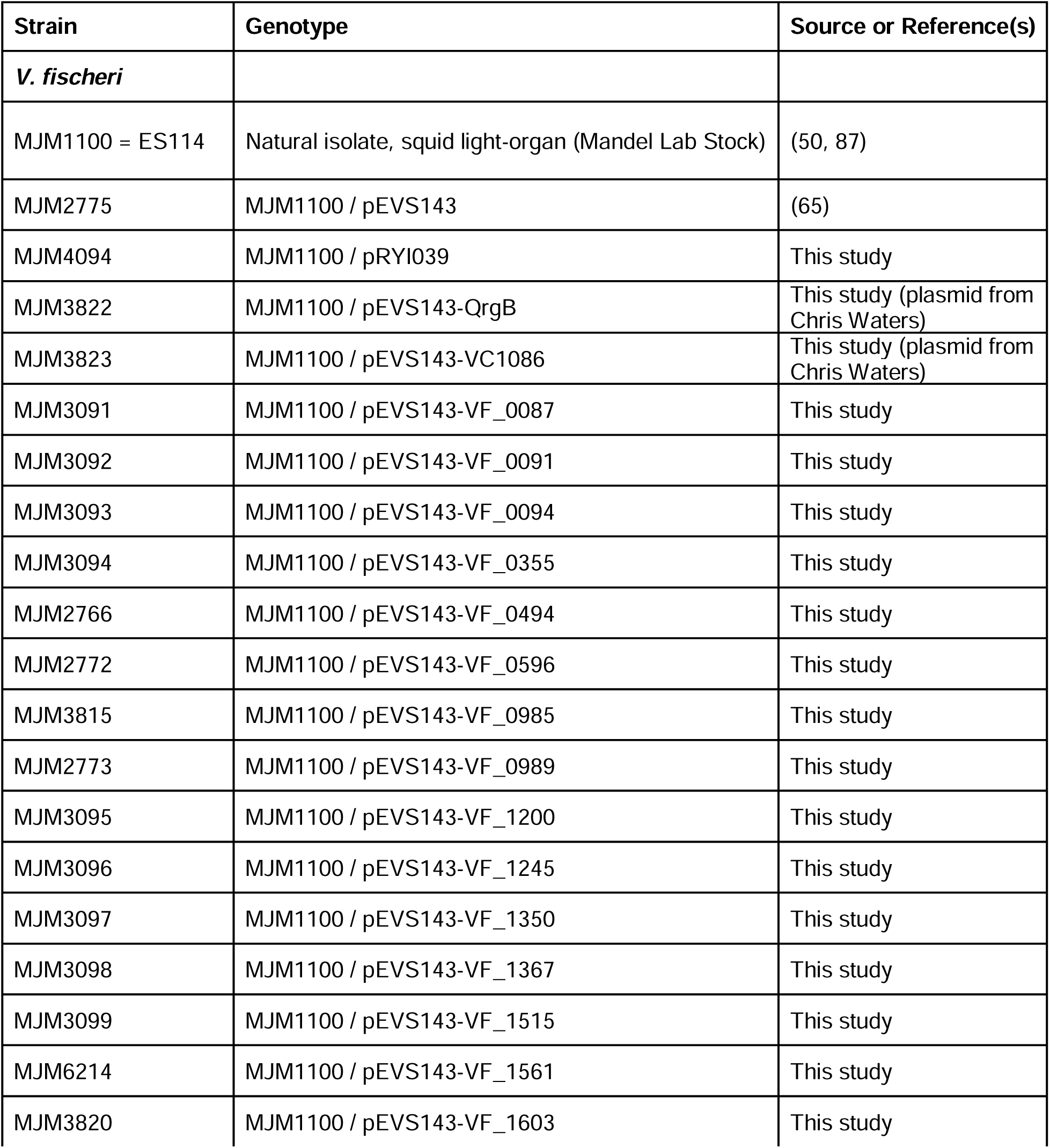
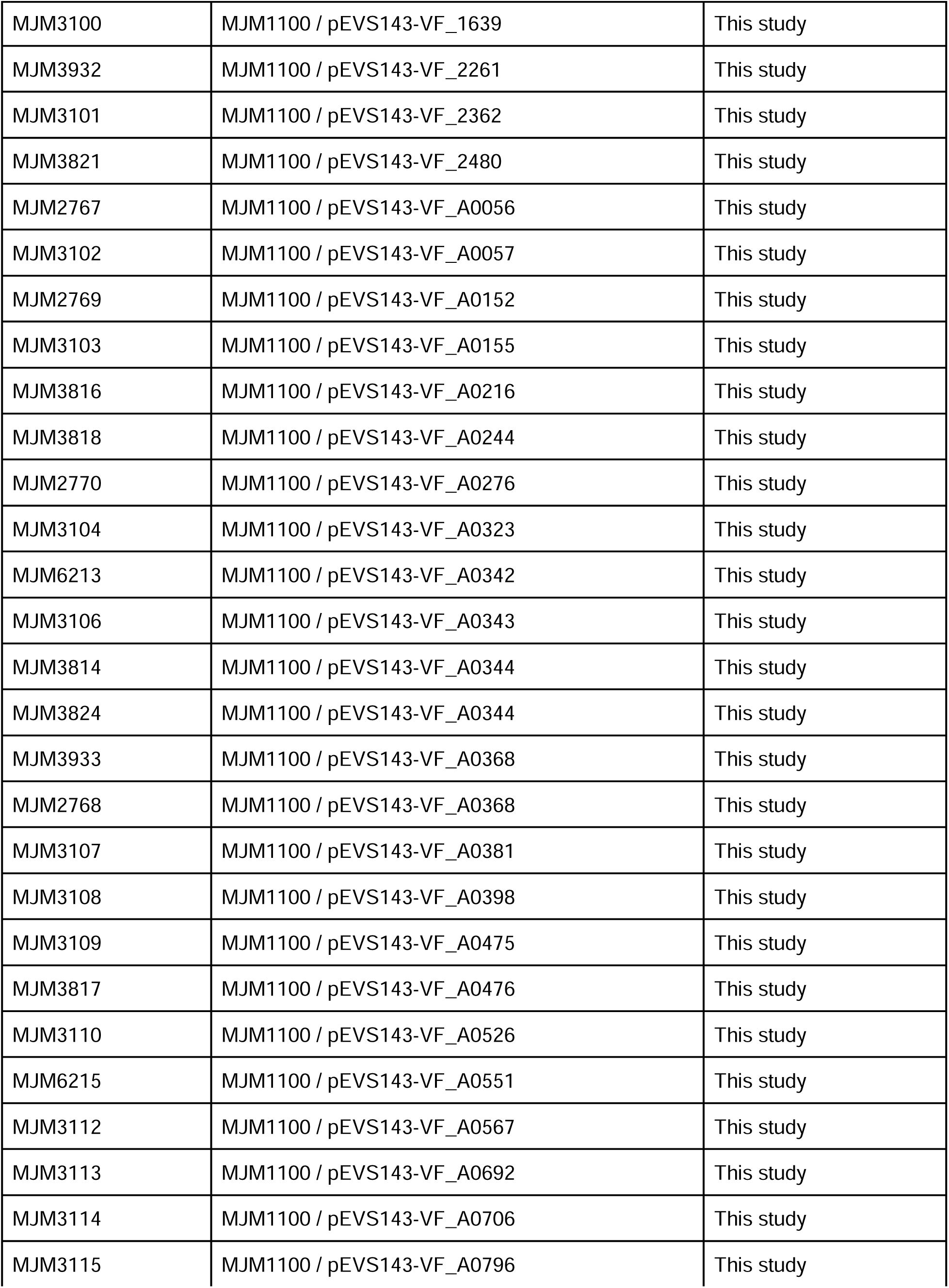
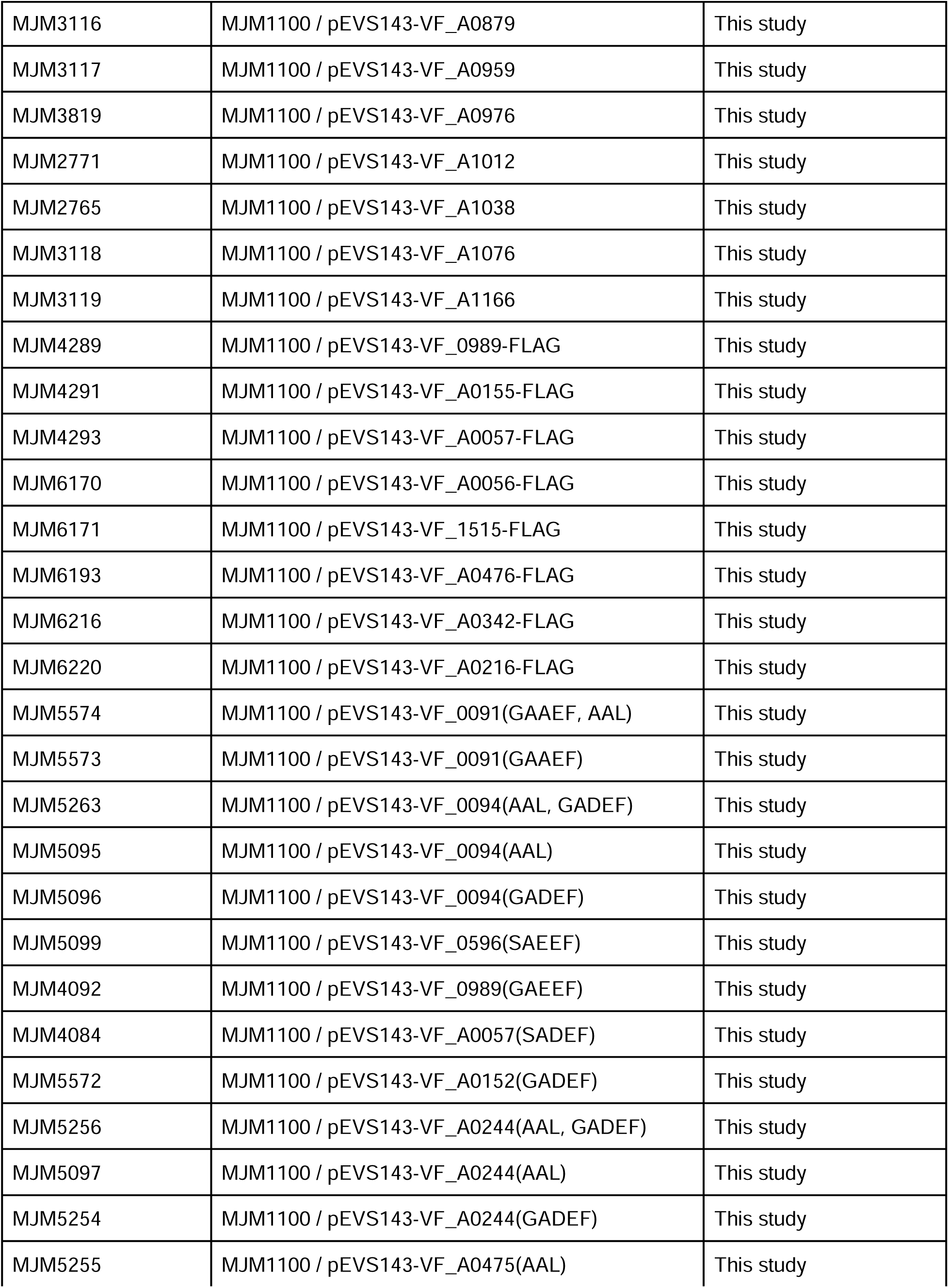
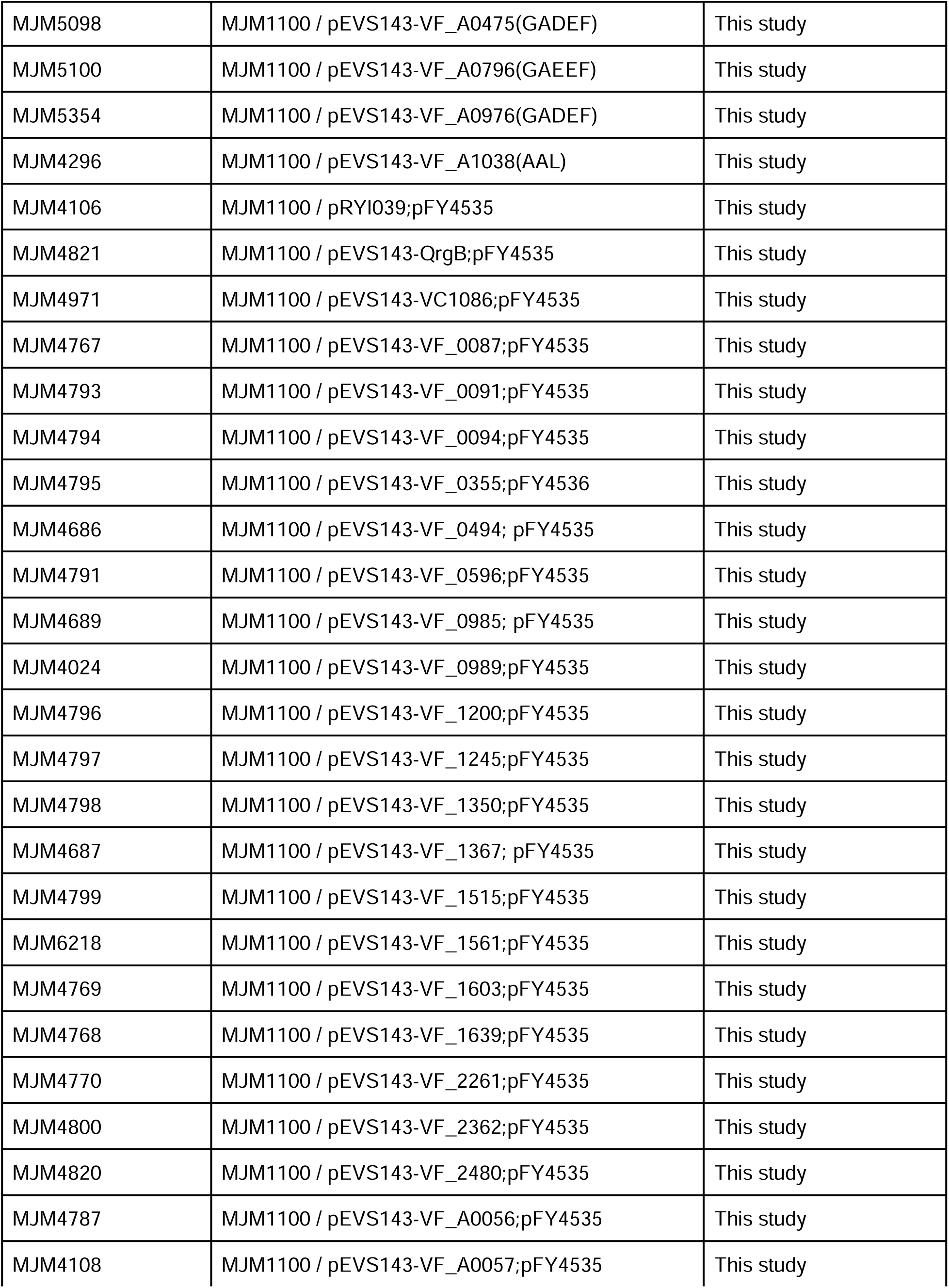
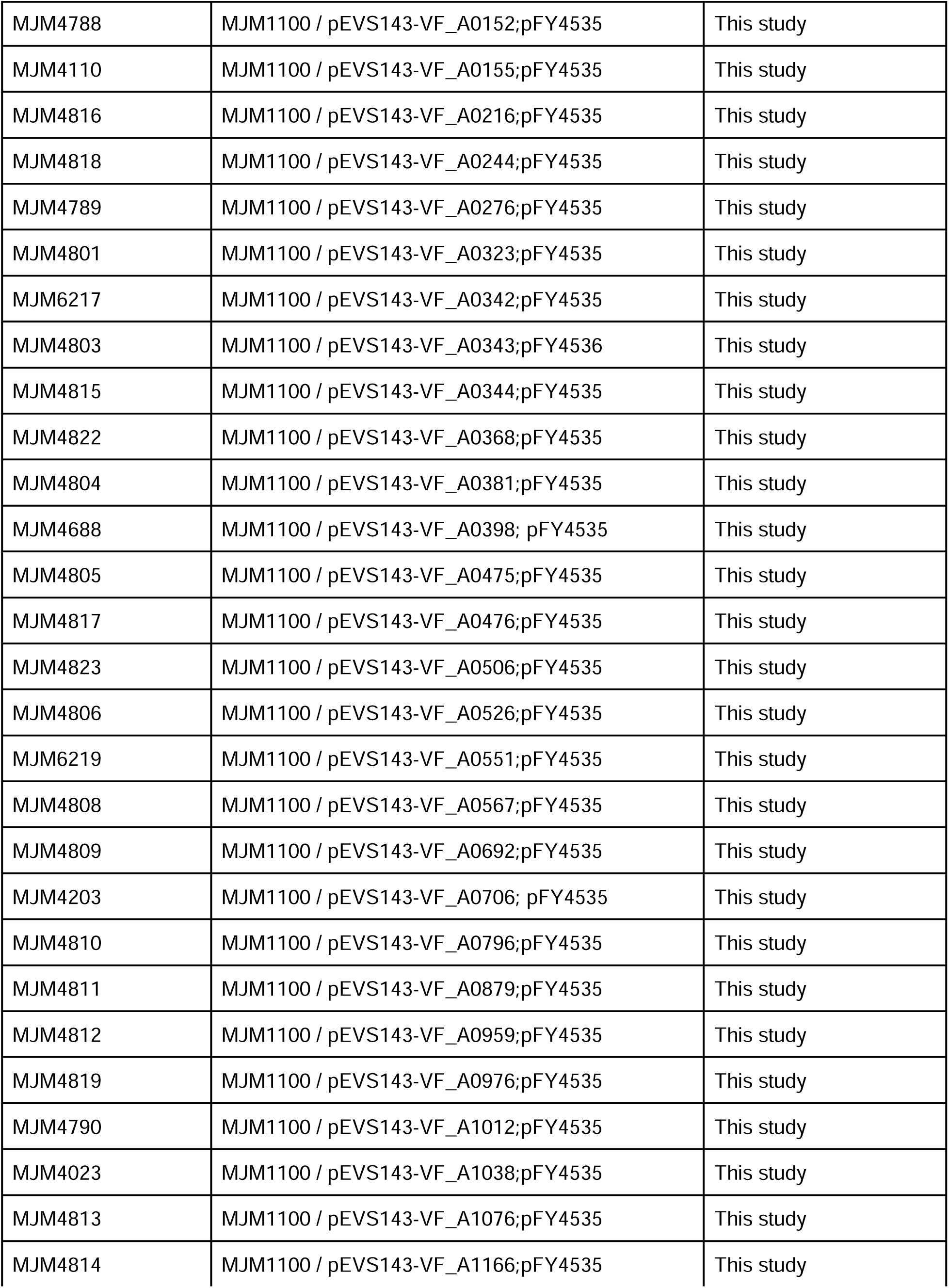
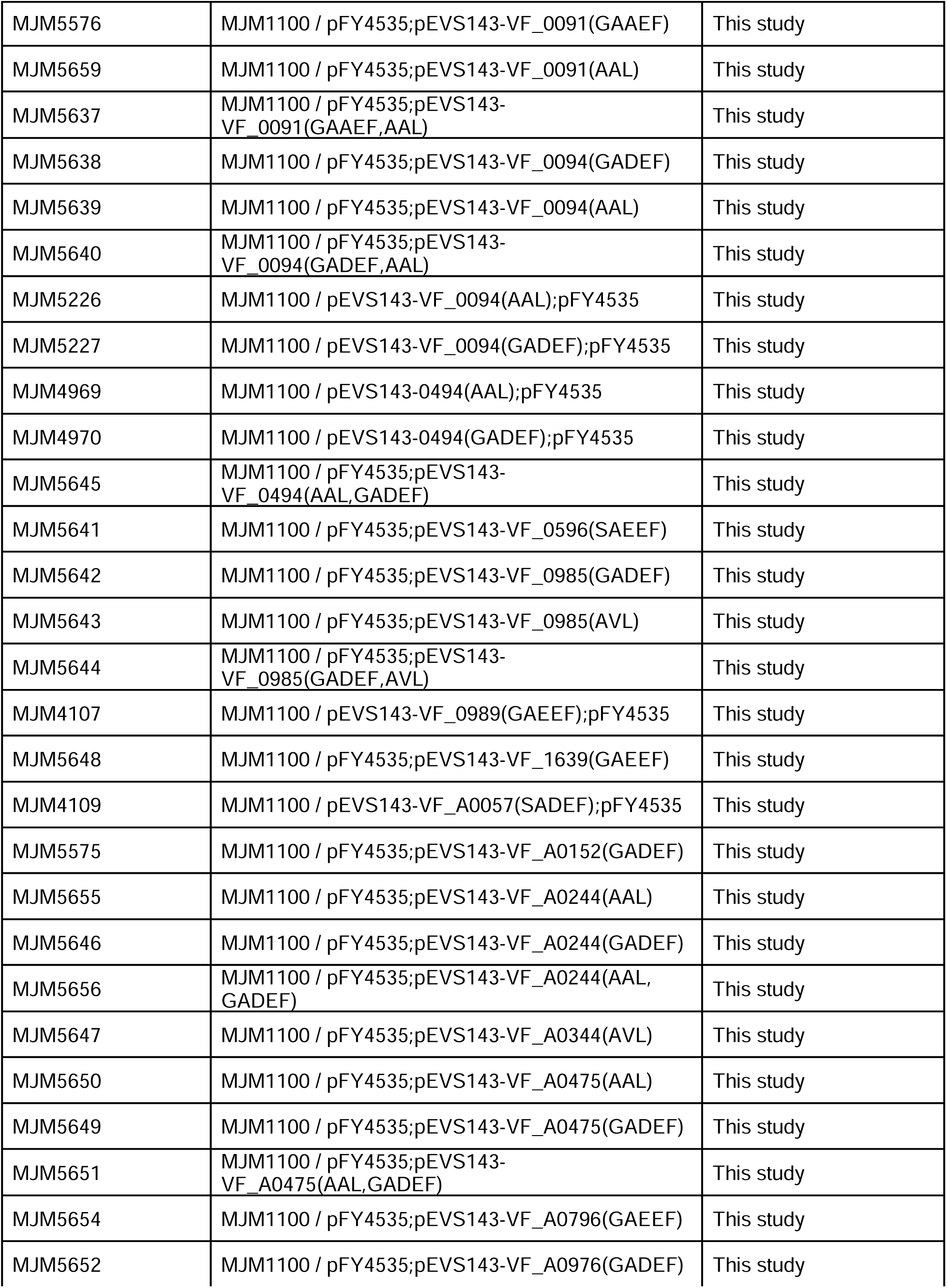
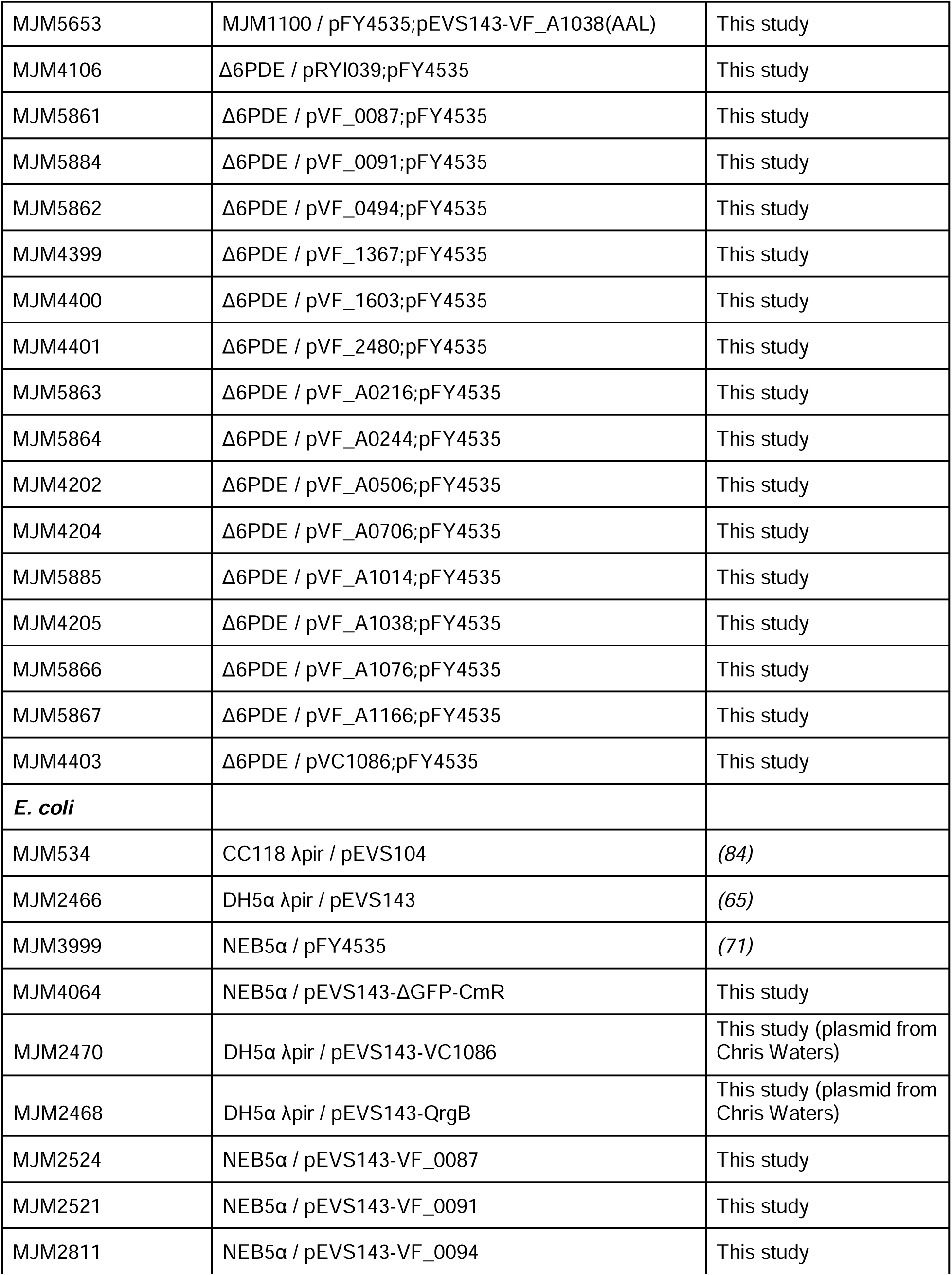
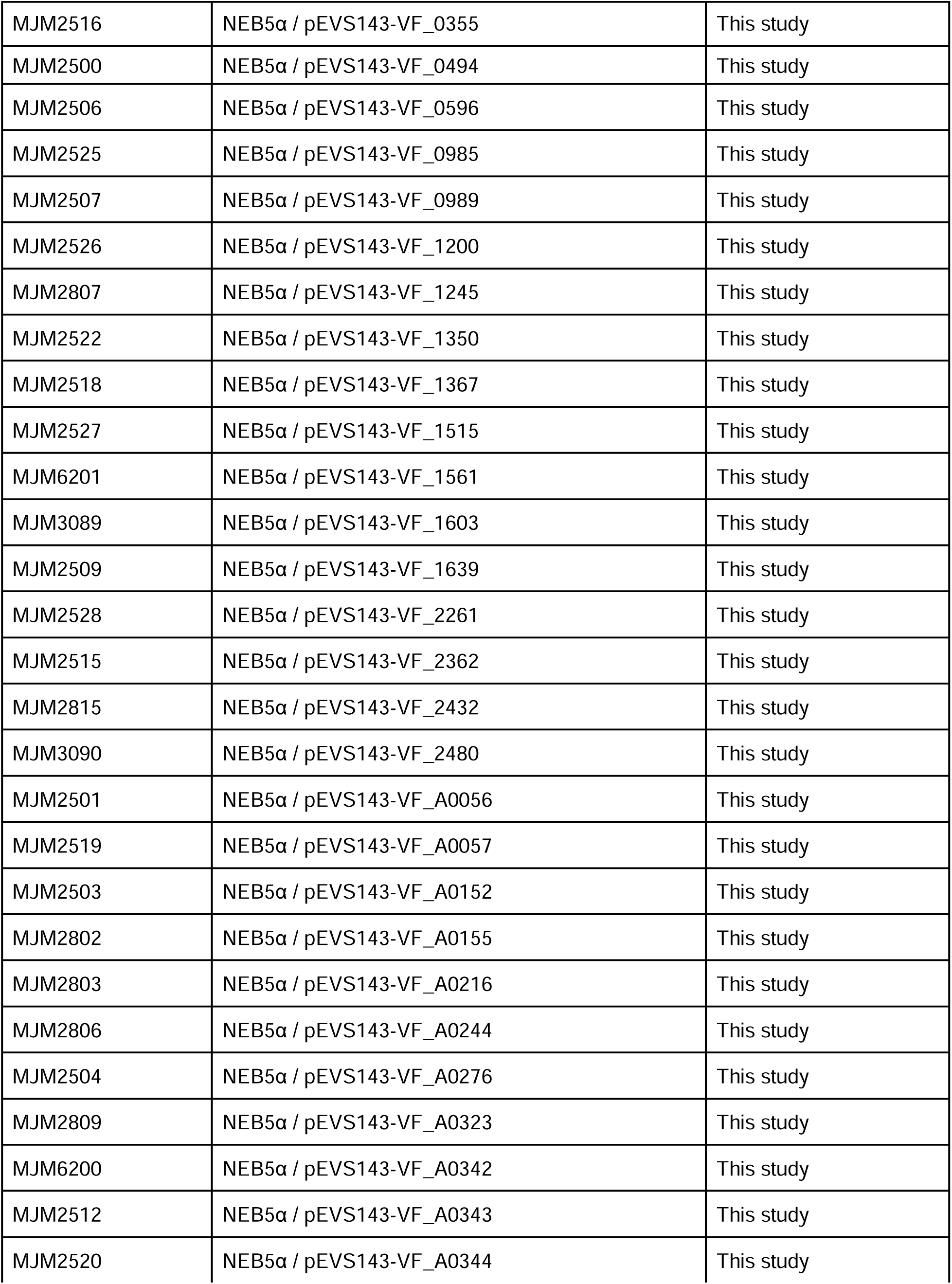
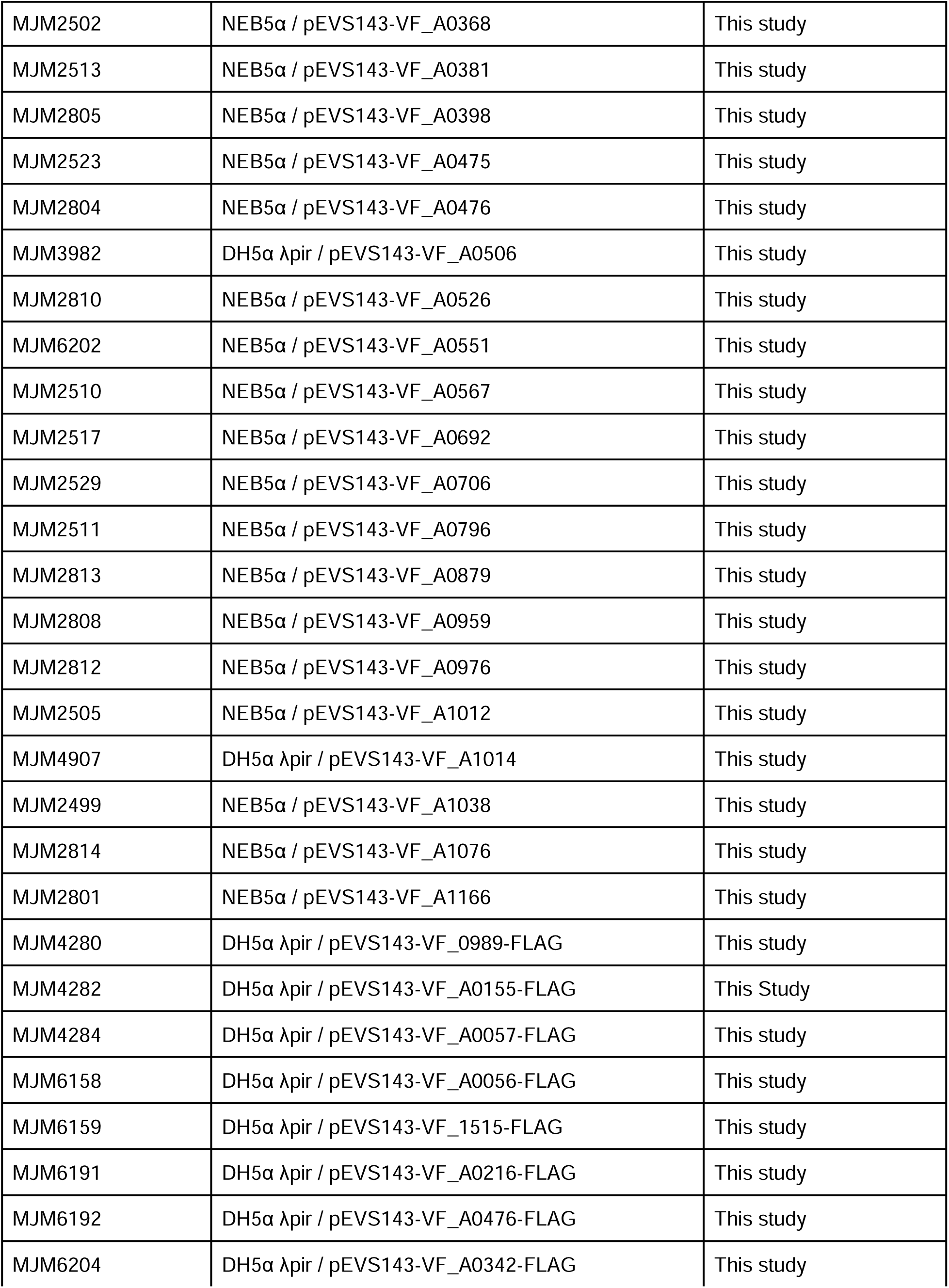
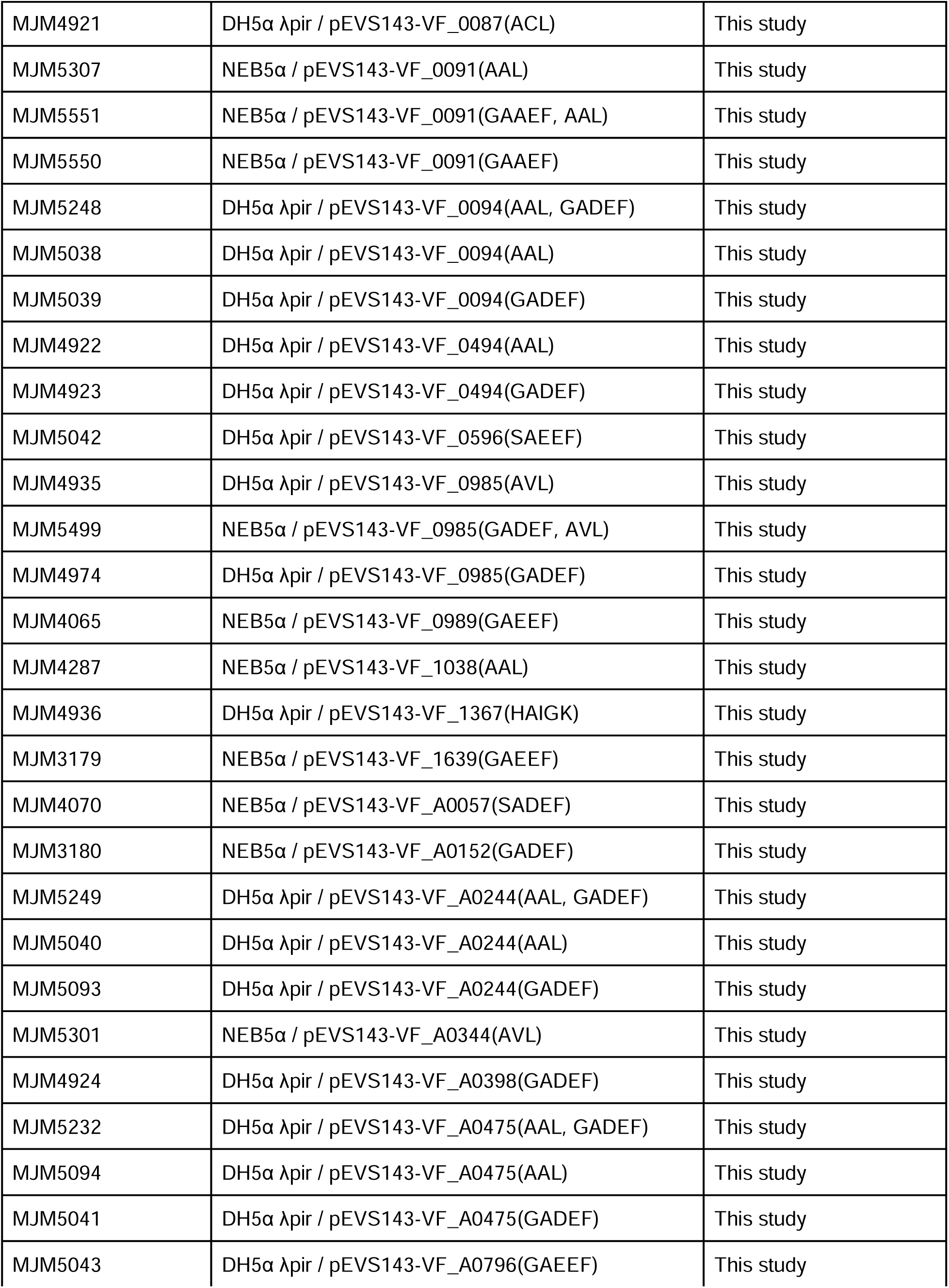
Strains.

**Table 2.**
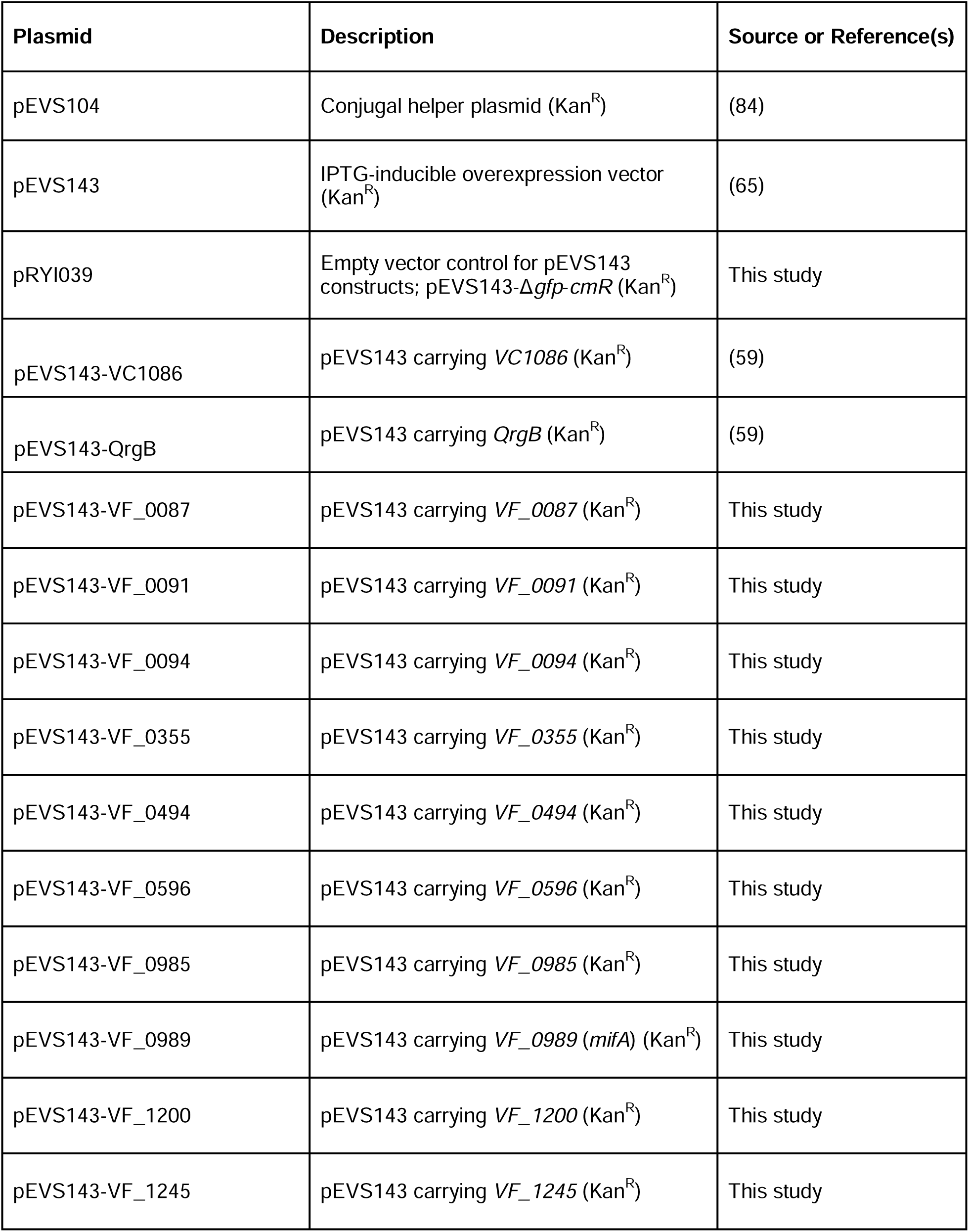
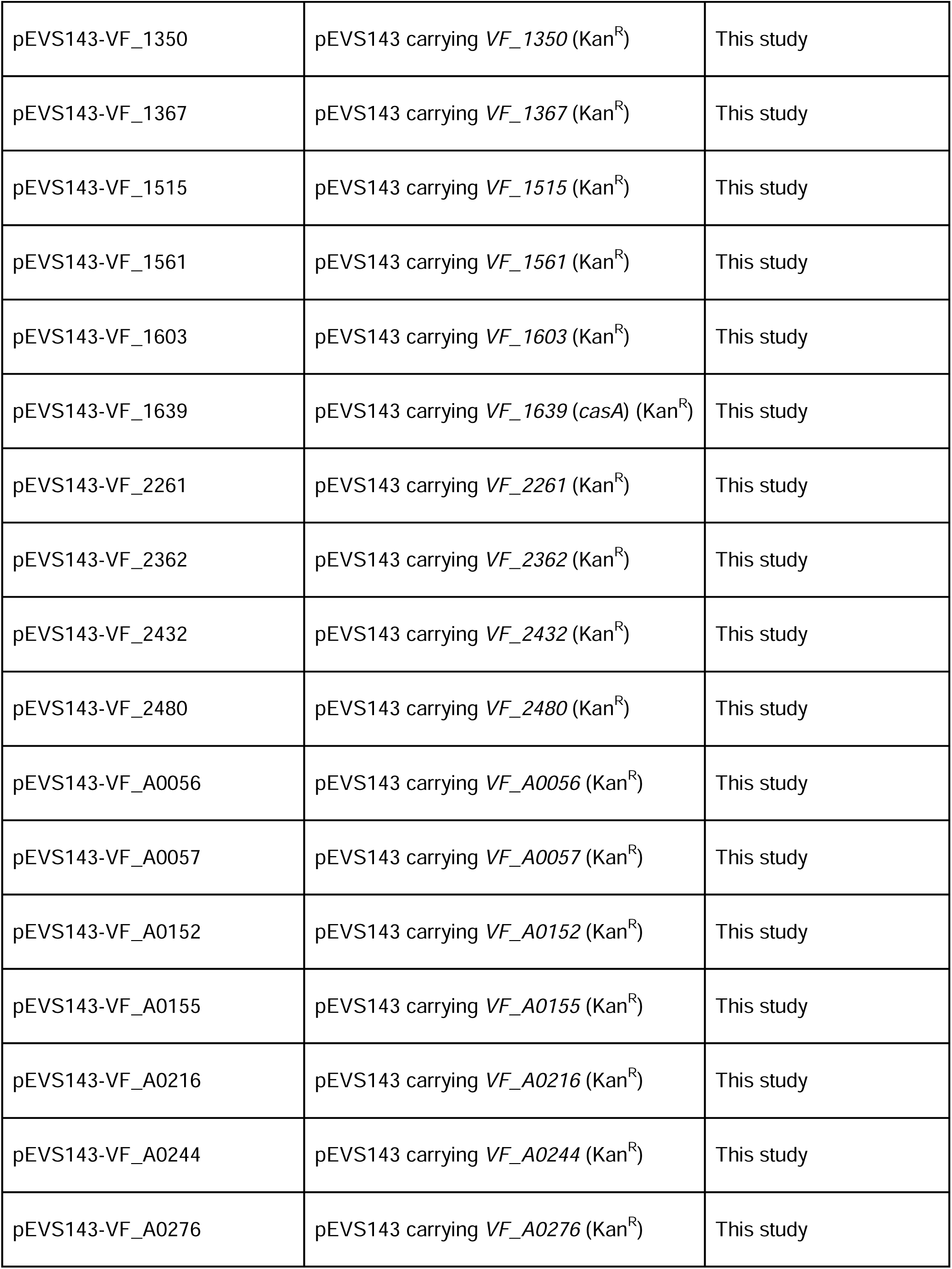
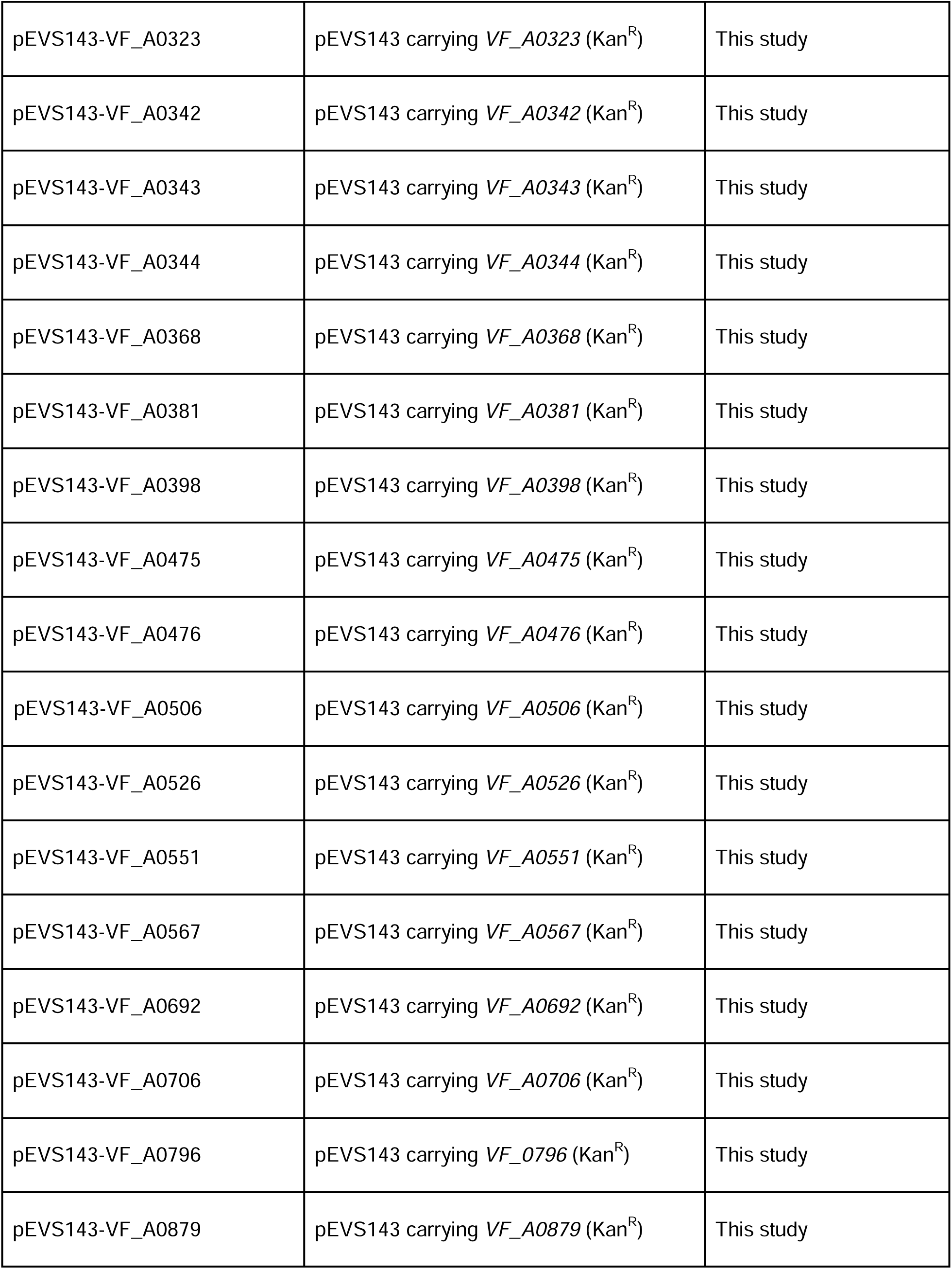
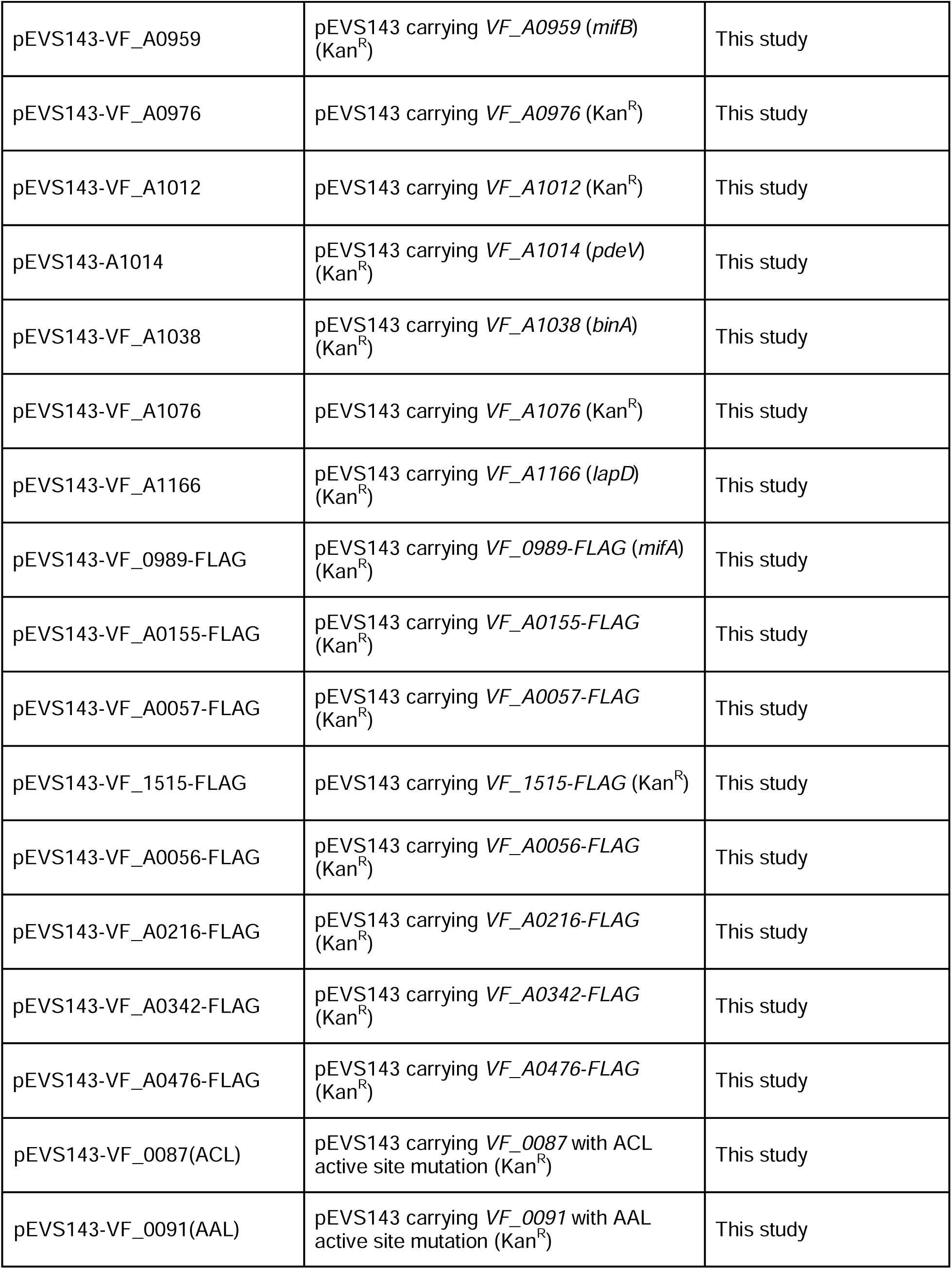
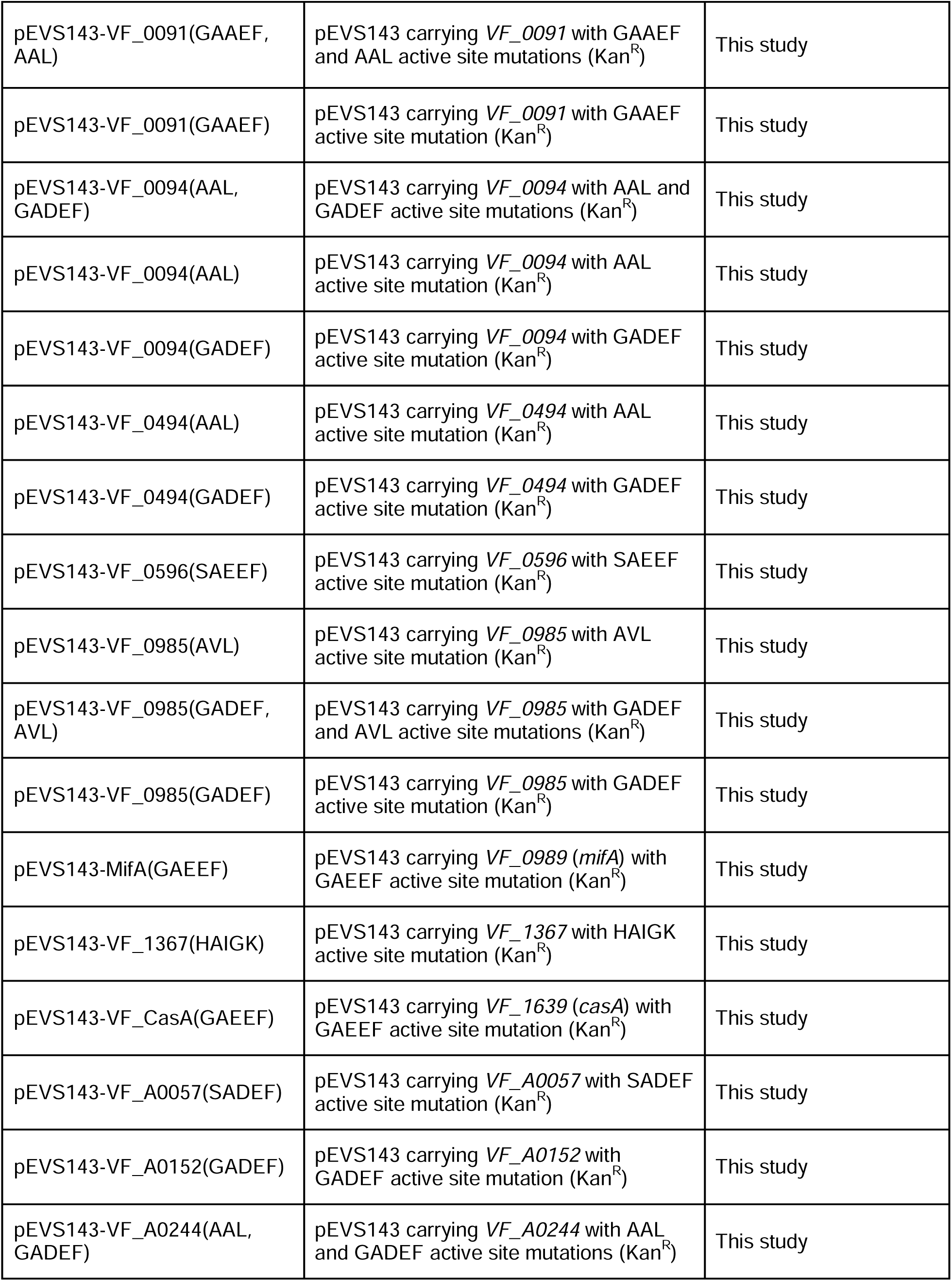
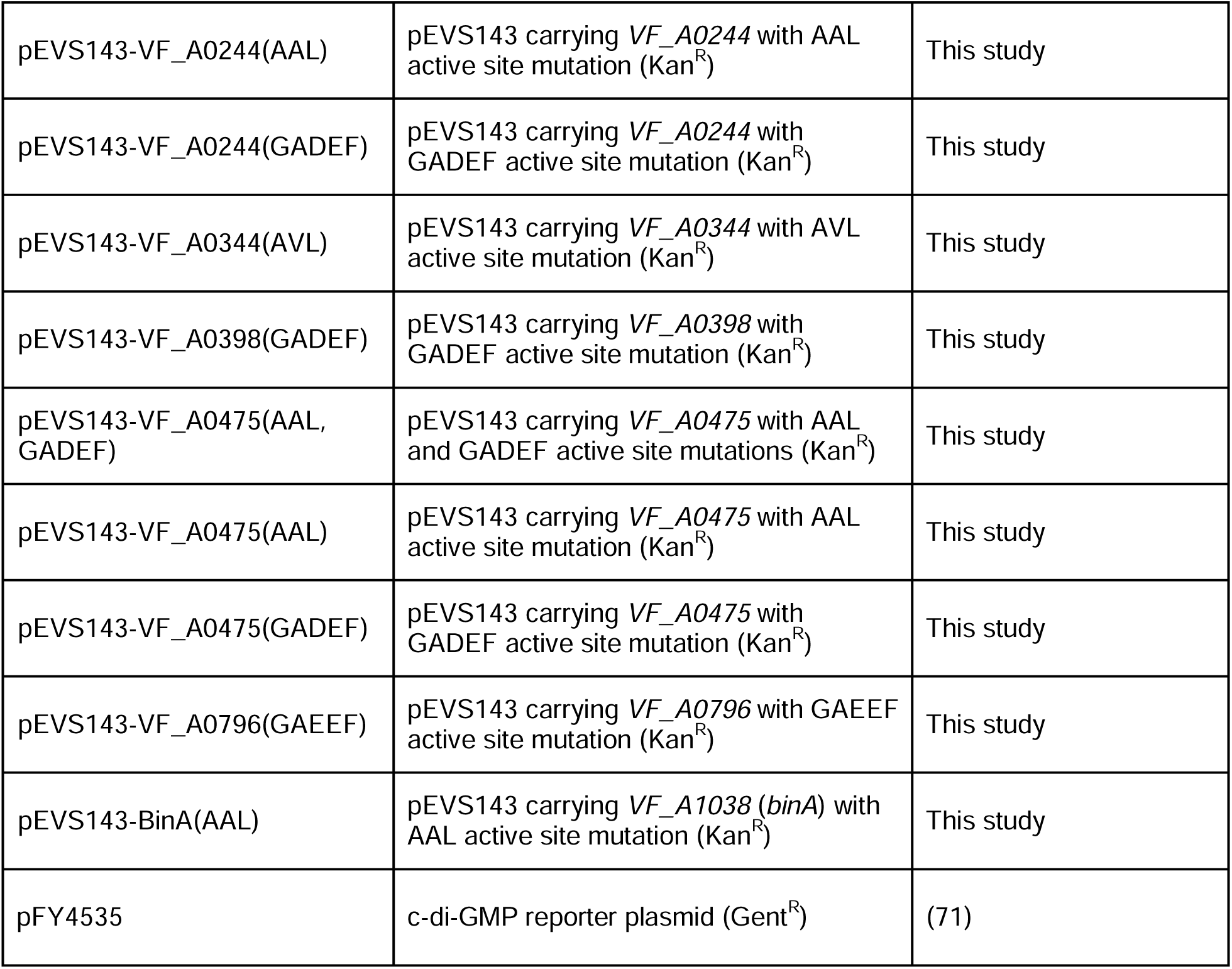
Plasmids.

### DNA synthesis and sequencing

Primers used in this study are listed in Table 3 and were synthesized by Integrated DNA Technologies (Coralville, IA). Gibson primers for all plasmids except pRYI039 and pEVS143-VF_A0506 were designed using the NEBuilder online tool. Site-directed mutagenesis primers were designed using the NEBaseChanger online tool. Full inserts for cloned constructs and active site mutant constructs were confirmed by Sanger Sequencing at Northwestern University Center for Genetic Medicine, ACGT, Inc. (Wheeling, IL), Functional Biosciences (Madison, WI), and/or whole plasmid sequencing by Plasmidsaurus (Eugene, OR). Sequence data were analyzed using DNASTAR or Benchling. PCR to amplify constructs for cloning and sequencing was performed using Pfx50 DNA Polymerase (Invitrogen) or Q5 High-Fidelity DNA polymerase (NEB). Diagnostic PCR was performed using GoTaq polymerase (Promega).

**Table 3.**
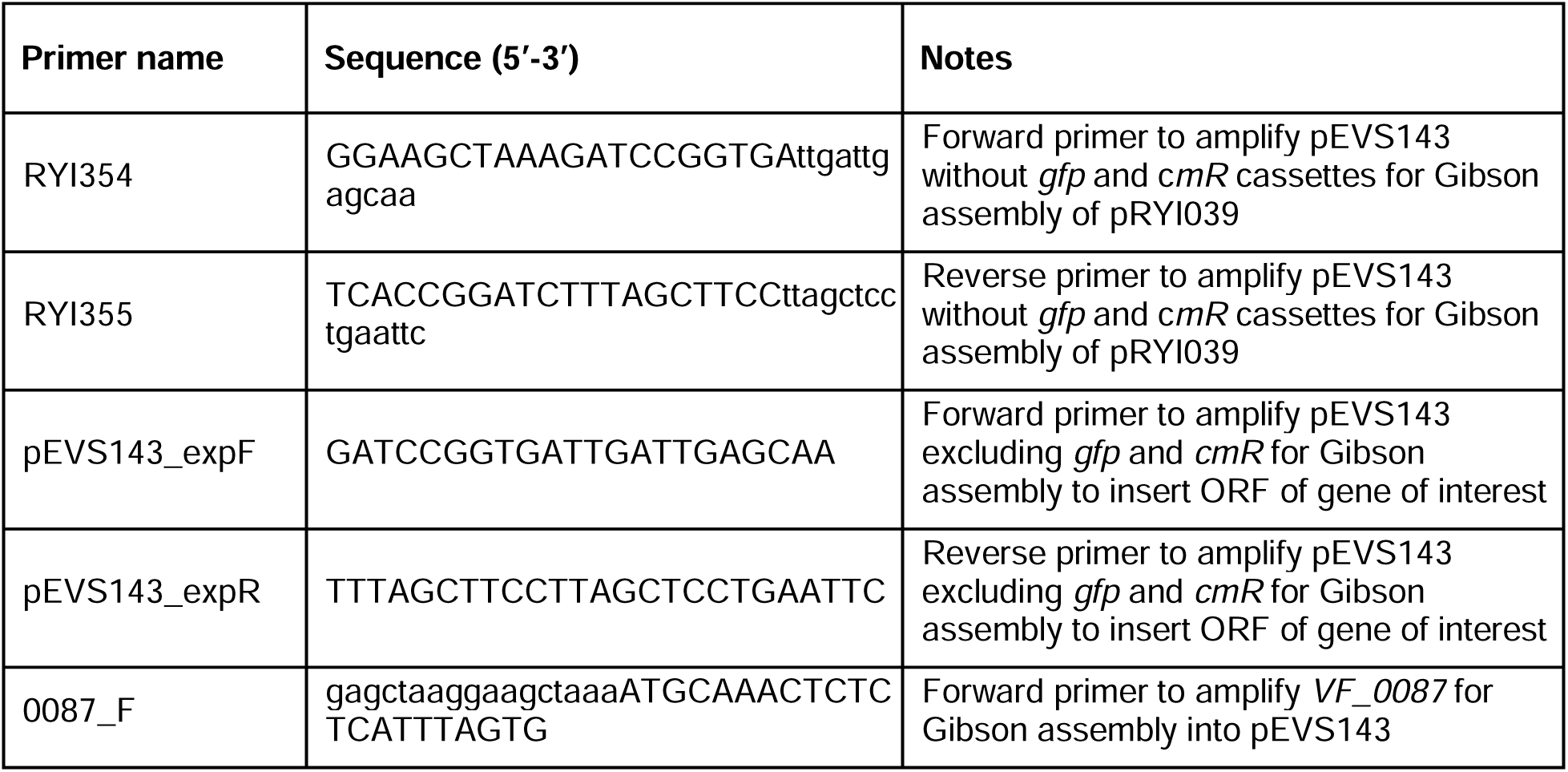
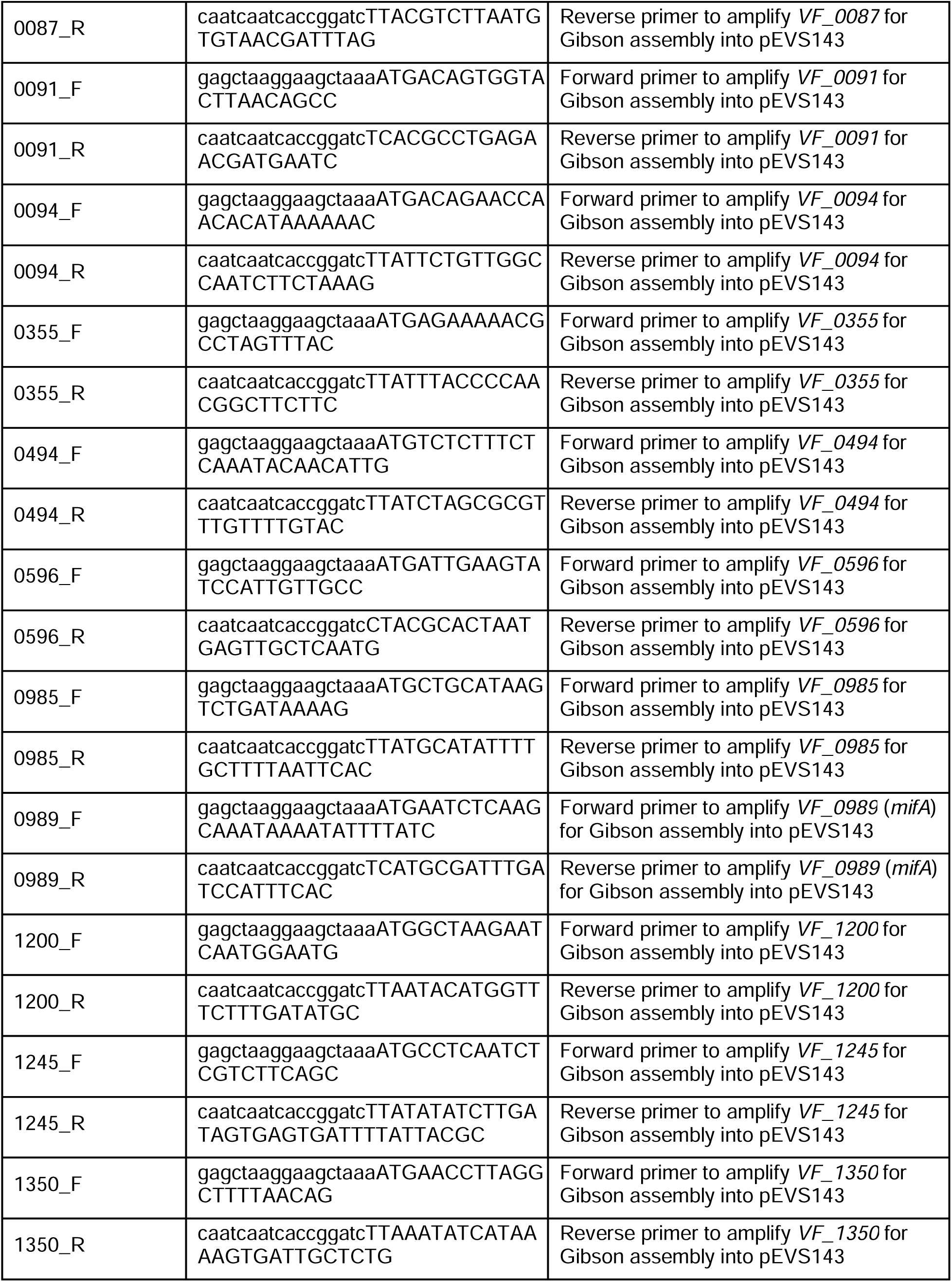
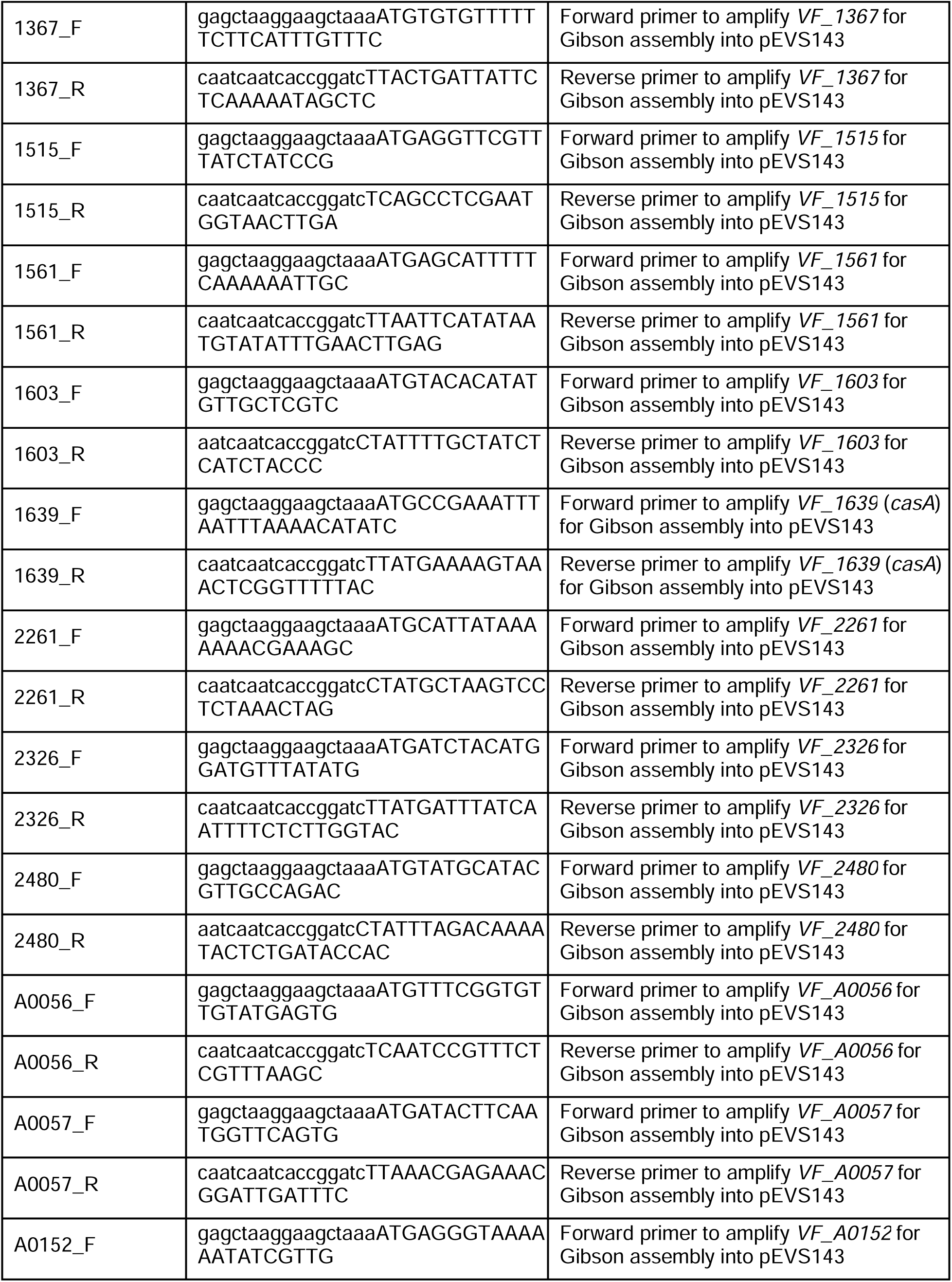
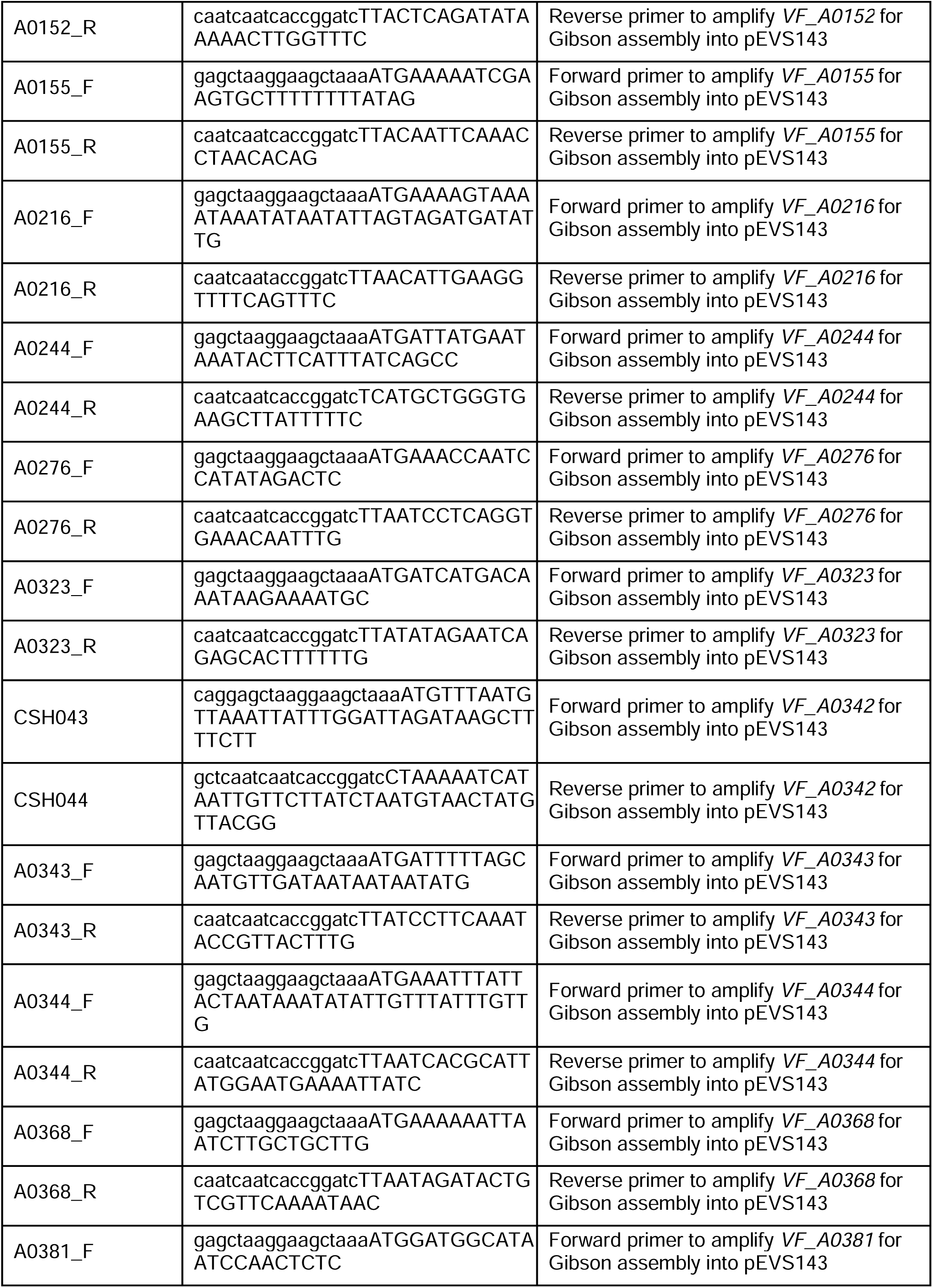
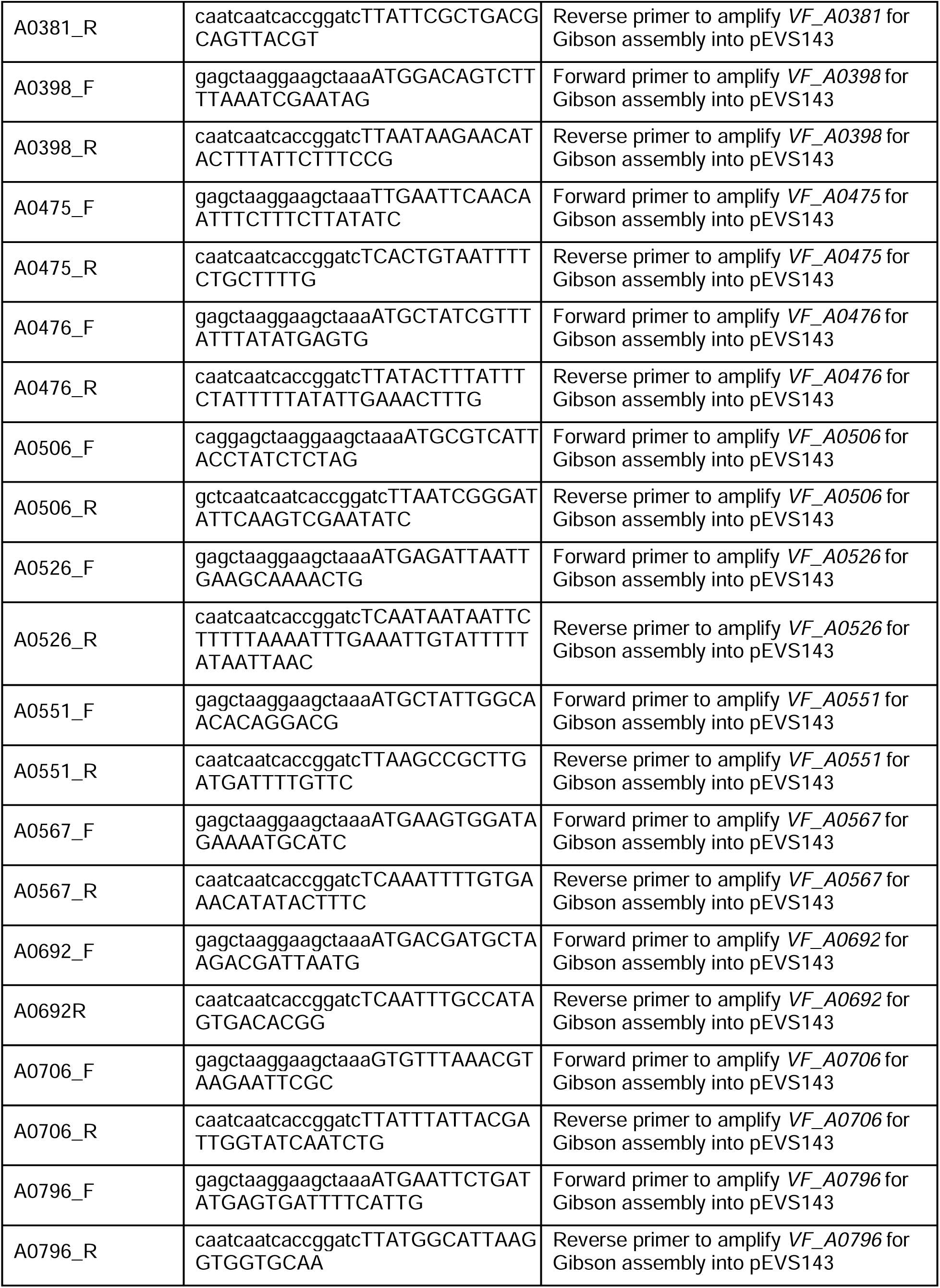
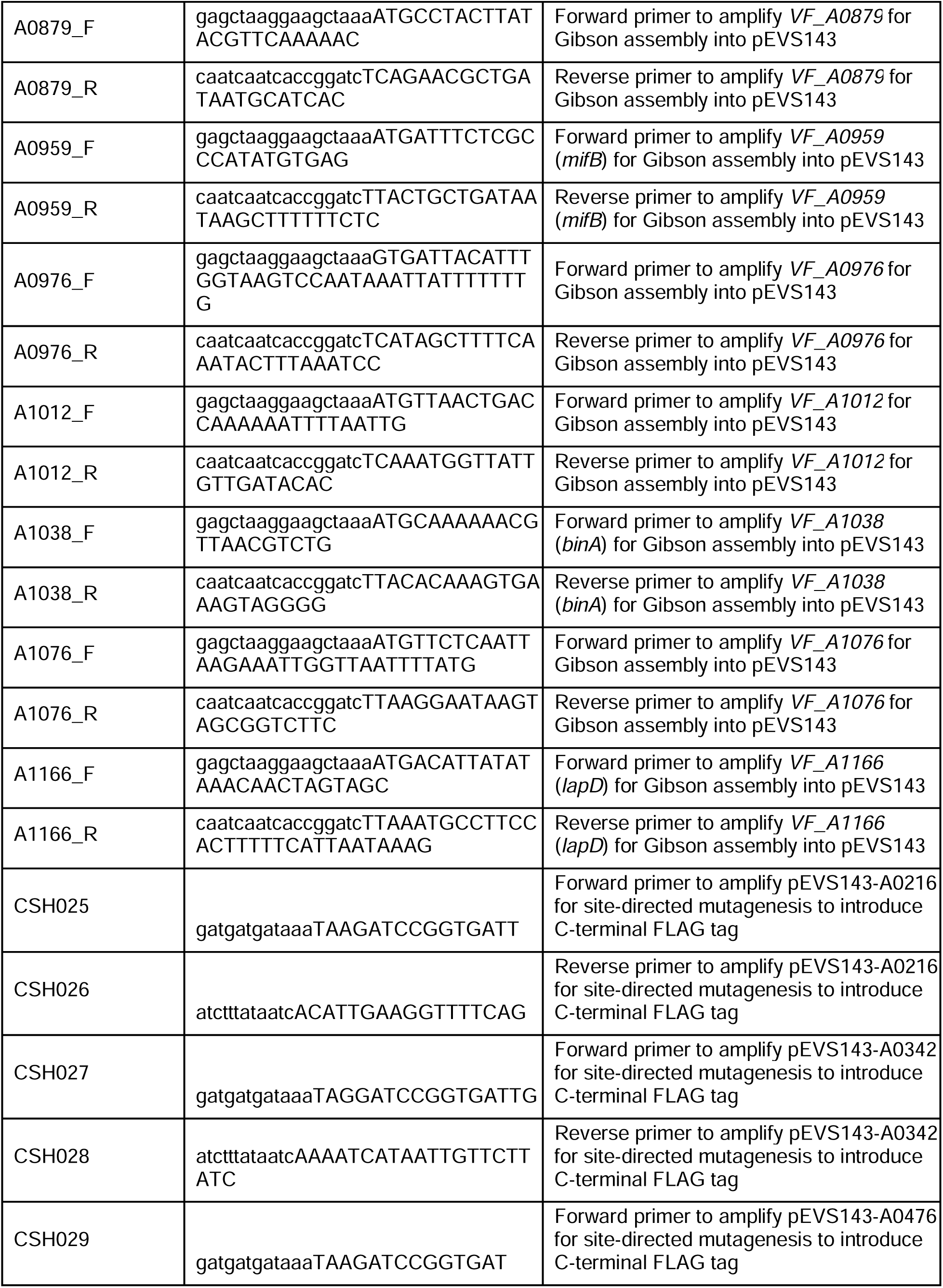
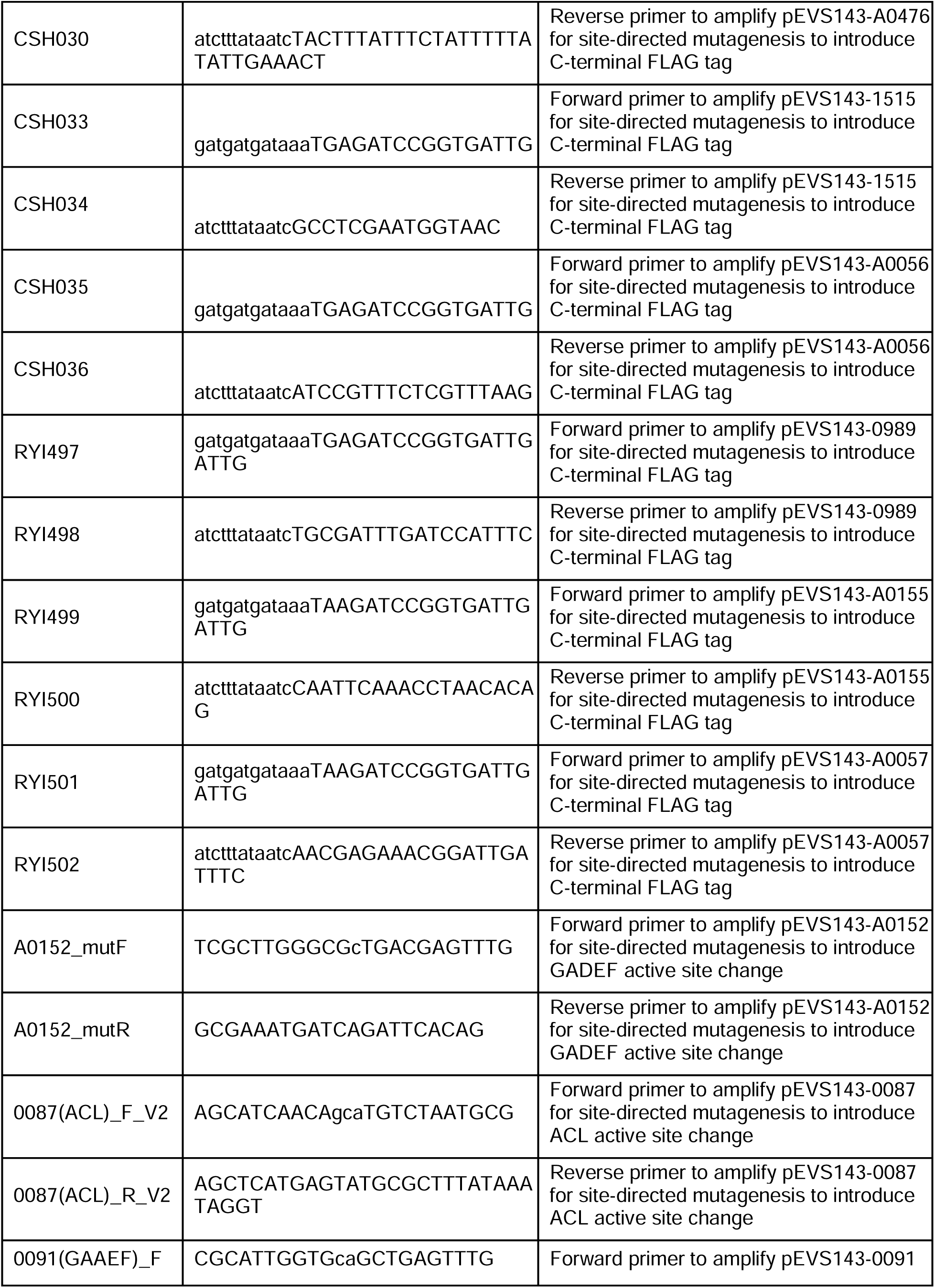
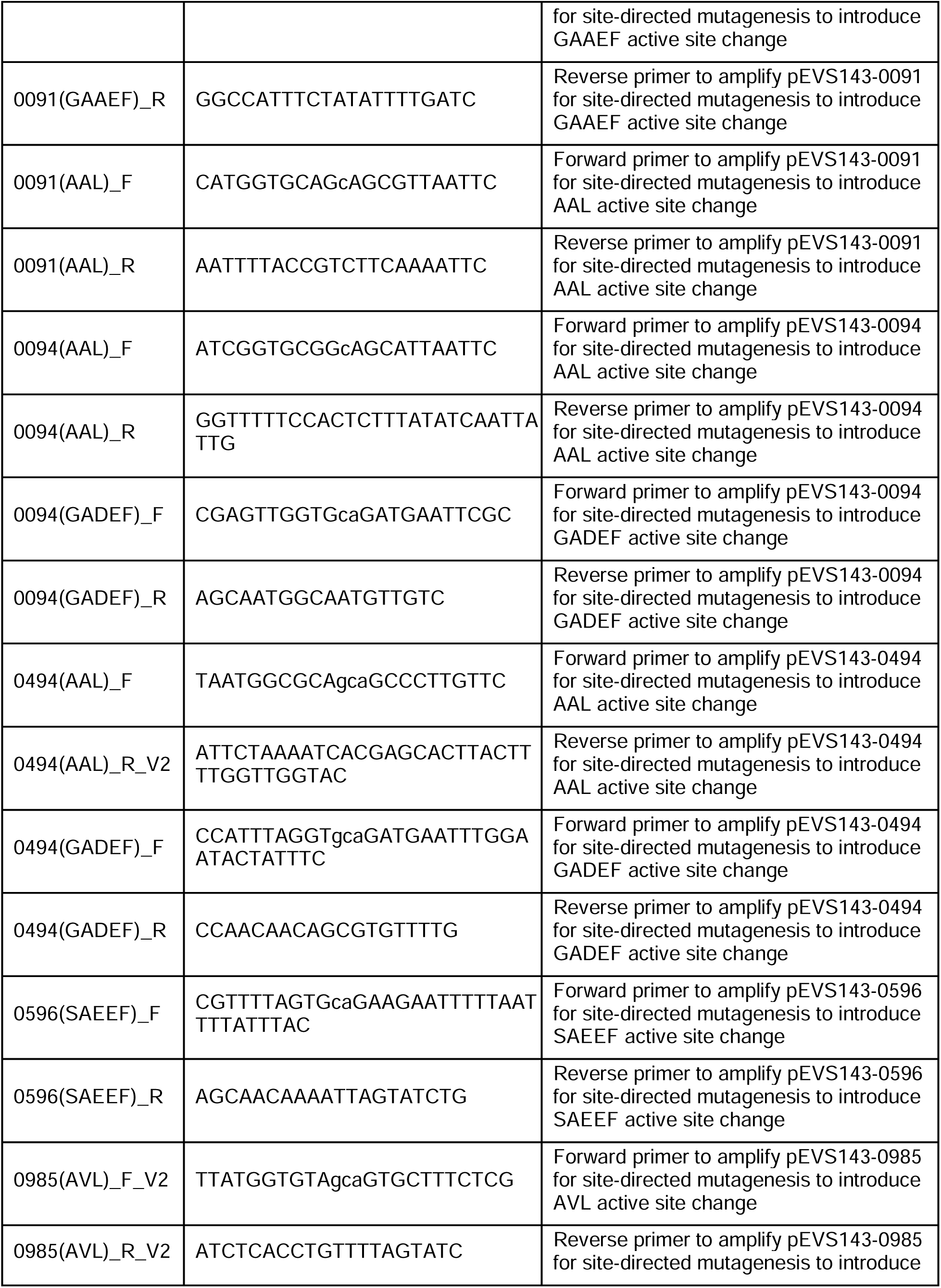
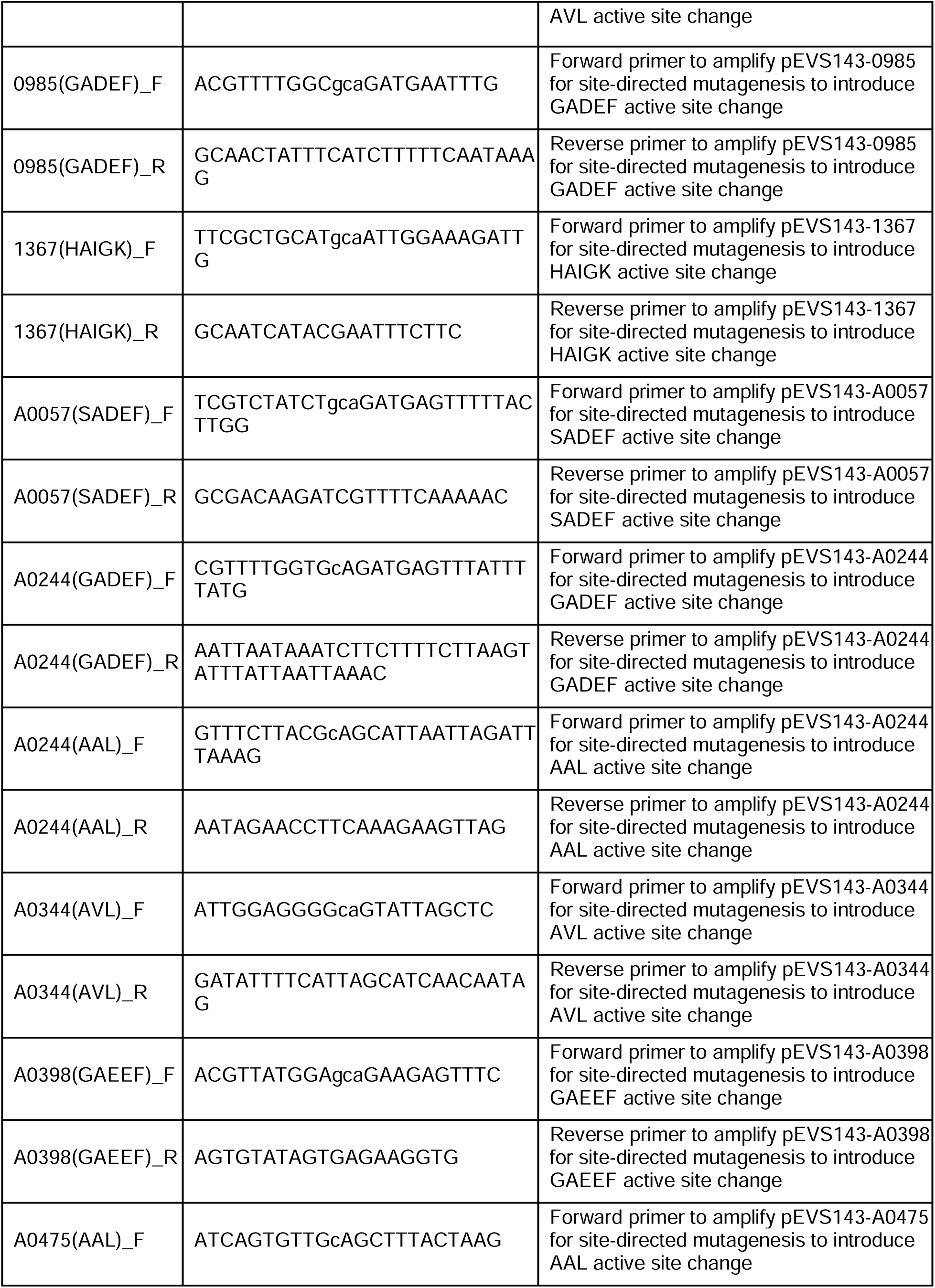
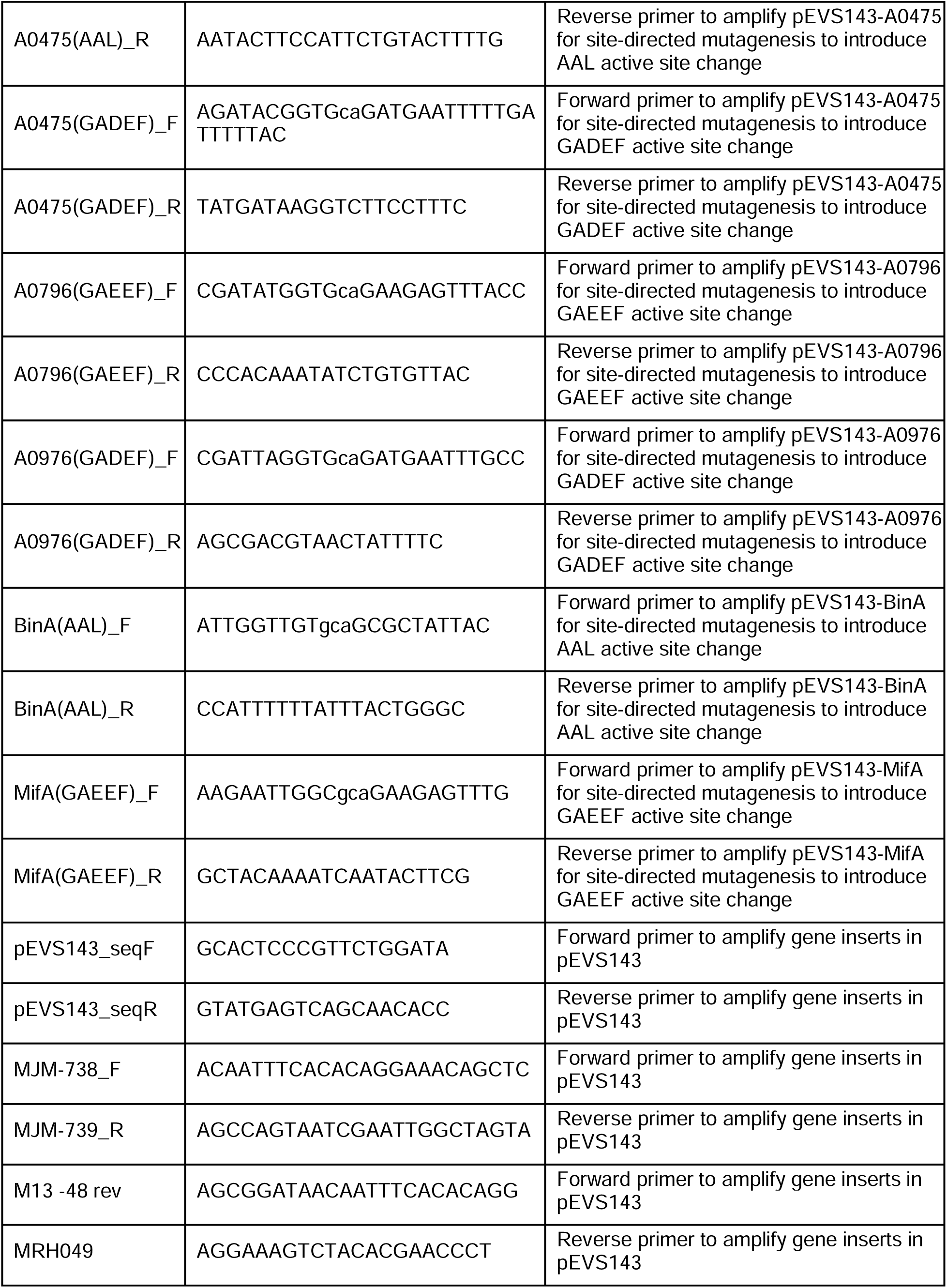
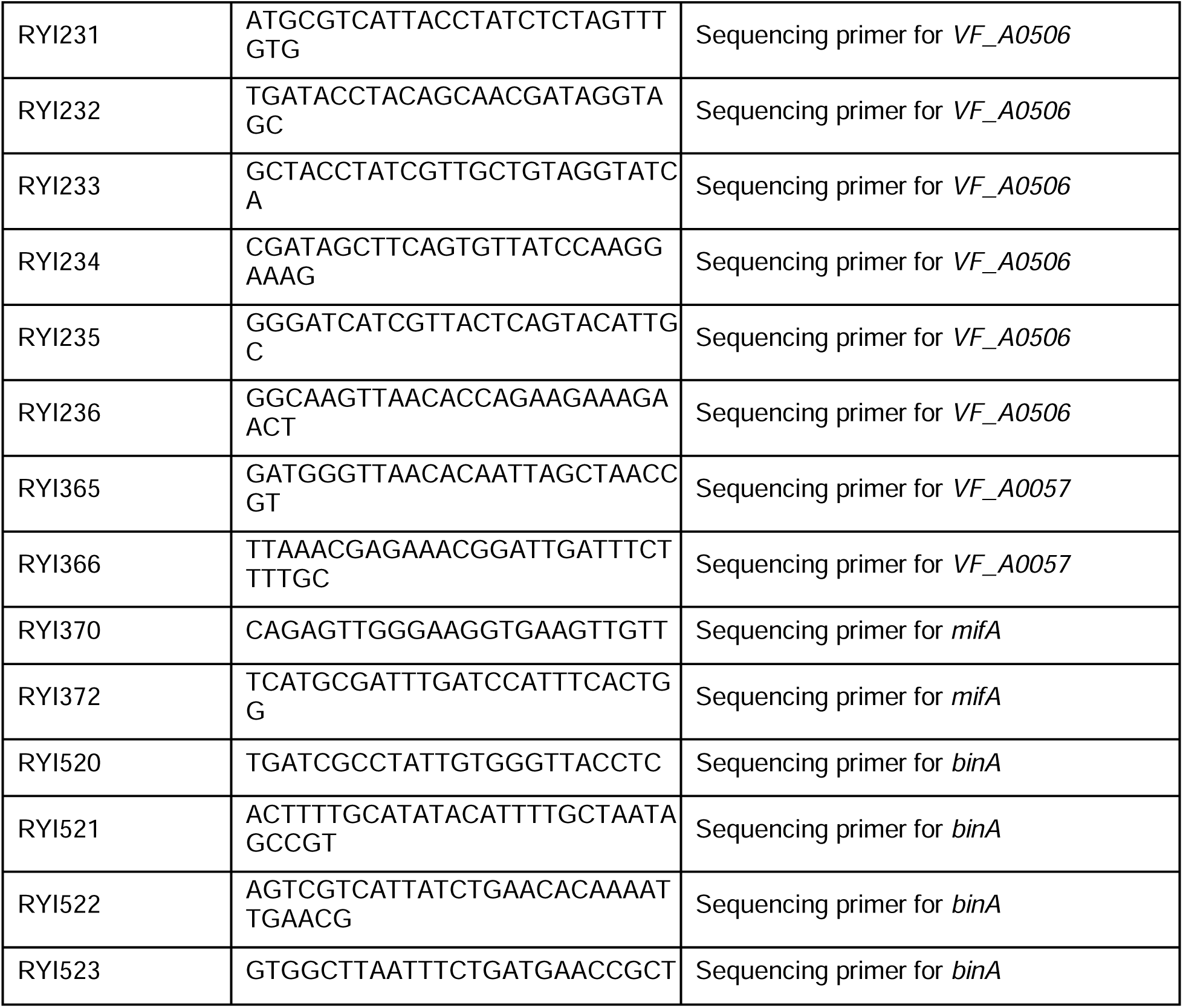
Primers.

### Construction of pRYI039

pEVS143 was amplified using primers RYI354 and RYI355. pRYI039 was assembled by Gibson assembly using the NEBuilder HiFi DNA Assembly Master Mix (NEB). The assembly reaction was transformed into chemically competent NEB5α cells and candidate transformants were selected using kanamycin. The plasmid was screened by PCR using primers M13 -48 rev and MRH049. The plasmid was confirmed by Sanger sequencing using primers M13 -48 rev and MRH049.

### Construction of the overexpression plasmids

pEVS143 was amplified using pEVS143_expF and pEVS143_expR primers. Gene ORFs were amplified from MJM1100 genomic DNA using gene-specific forward and reverse primers. Amplified gene inserts were assembled with amplified pEVS143 by Gibson assembly using NEB Gibson Assembly Cloning Kit (NEB) or NEBuilder HiFi DNA Assembly Master Mix (NEB). The assembly reactions were transformed into chemically competent NEB5α or DH5α λpir cells and candidate transformants were selected using kanamycin. Each plasmid was screened by PCR using primers MJM-738F and MJM739R; M13 -48 rev and MJM-739R; or MJM-738F and the respective assembly reverse primer, MJM-739R and the assembly forward primer, and/or MJM-738F and MJM739R. To construct pEVS143-VF_A1014, *VF_A1014* was synthesized with flanking AvrII and BamHI restriction sites and cloned into pEVS143 by GenScript (Piscataway, NJ) and plasmid was confirmed by whole plasmid sequencing. pEVS143-VF_A0506 was confirmed by Sanger sequencing using primers M13 -48 rev, RYI225, RYI226, RYI227, RYI228, RYI229, RYI230, and MRH049. Remaining plasmids were confirmed by Sanger sequencing using primers MJM-738F and MJM-739R; MJM-738F and the respective assembly reverse primer and/or MJM-739R and the assembly forward primer; or pEVS143_seqF and pEVS143_seqR.

### Construction of FLAG-tagged protein overexpression plasmids

Plasmids were amplified using site-directed mutagenesis primers to introduce active site mutations. Active site mutant plasmids were assembled using the Q5 Site-Directed Mutagenesis Kit (NEB). Site-directed mutagenesis reactions were transformed into chemically competent DH5α λpir cells and candidate transformants were selected using kanamycin. Each plasmid was screened by PCR using primers M13 -48 rev and MJM-739R and confirmed by whole plasmid sequencing.

### Construction of the DGC and PDE active site mutant overexpression plasmids

Plasmids were amplified using site-directed mutagenesis primers to introduce active site mutations. Active site mutant plasmids were assembled using the Q5 Site-Directed Mutagenesis Kit (NEB). Site-directed mutagenesis reactions were transformed into chemically competent NEB5α or DH5α λpir cells and candidate transformants were selected using kanamycin. Each plasmid was screened by PCR using primers M13 -48 rev and MJM739R; or MJM-738F and the respective assembly reverse primer, MJM-739R and the assembly forward primer, and MJM-738F and MJM739R. pEVS143-VF_0989(GAEEF) was confirmed by Sanger sequencing using primers RYI370 and RYI372. pEVS143-VF_A0057(SADEF) was confirmed by Sanger sequencing using primers RYI365 and RYI366. pEVS143-VF_A1038 was confirmed by Sanger sequencing using primers RYI520, RYI521, RYI522, RYI523, MJM-738F, and MJM-739R. Remaining active site mutant plasmids were confirmed by whole plasmid sequencing or Sanger sequencing using primers MJM-738F and MJM-739R; MJM-738F and the respective template plasmid Gibson assembly reverse primer and/or MJM-739R and the template plasmid Gibson assembly forward primer; or pEVS143_seqF and pEVS143_seqR.

### Assembly of arrayed strain collections

Strains were streaked on LBS agar and incubated overnight at 25°C. Liquid LBS was inoculated with single colonies of each strain in a deep-well 96-well plate and grown overnight at 23-25°C on a shaker. Overnight cultures were saved in glycerol stocks in 96-well plates in triplicate. For VF_1561, VF_A0152, VF_A0342, and VF_A0551, errors were found in the strains in the wild-type background in the original assays, and the corrected strains were reanalyzed in the same 96-well format in a new arrayed strain collection alongside select additional strains (pRYI039, pEVS143, VF_0087, and VF_0091, VF_0985, and MifA) as controls.

### Congo red biofilm assay

Liquid LBS was inoculated with 2 μL of glycerol stock of each strain from an arrayed strain collection and grown at room temperature (23-25°C) overnight on a shaker. 2 μL spots of liquid culture were spotted on LBS Congo red agar and incubated 24 h at 25°C. Spots were transferred onto white printer paper (86) and images were scanned as TIFF or JPEG files. Congo red binding was quantified using ImageJ software (version 2.0.0-rc-69/1.52p) as described in the “Data analysis” section below.

### Motility assay

Liquid LBS was inoculated 2 μL of glycerol stock of each strain from an arrayed strain collection and grown at room temperature (23-25°C) overnight on a shaker. 2 μL spots of liquid culture were spotted on TBS, TBS-Mg^2+^ (35 mM MgSO_4_), and TBS-Ca^2+^ (10 mM CaCl_2_) agar and incubated overnight at 25°C. The spotted strains were inoculated into TBS, TBS-Mg^2+^, and TBS-Ca^2+^ soft (0.3%) agar, respectively, using a 96-pin replicator and incubated at 25°C for 3 h for TBS plates and 2.5 h for TBS-Mg^2+^ and TBS-Ca^2+^ plates. Images of plates were taken using a Nikon D810 digital camera and diameter of migration was measured using ImageJ software (version 2.0.0-rc-69/1.52p).

### Western blot analysis

One milliliter of overnight culture was pelleted then washed and lysed in 1% SDS solution. To standardize the total protein concentration, the volume of SDS was adjusted based on the Optical Density at 600 nm (OD_600_). Lysed cells were pelleted to remove cell debris, and the solution was mixed at a 1:1 ratio with 2x Laemmli sample buffer from Bio-Rad (Hercules, CA) to which β-mercaptoethanol had been added. Solution was heated at 95°C for 15 min and loaded onto a 4-20% Mini-Protean TGX Precast Stain Free Gel from Bio-Rad (Hercules, CA). Gel was transferred to a Immun-Blot Low Fluorescence polyvinylidene difluoride (PVDF) membrane and blocked overnight in 5% nonfat milk resuspended in 1x Tris-buffered saline Tween-20 (1x TBS-T). 2 µL of anti-FLAG Rabbit IgG (800 µG/mL, Sigma-Aldrich) was used as the primary antibody in 0.5% nonfat milk suspended in 1x TBS-T. 2 µL of anti-RpoA Mouse IgG (500 µG/mL, Biolegend) was used as a loading control, binding the RNAP α subunit. 2 µL of LI-COR IRDye 800CW Goat anti-Rabbit IgG (1,000 µg/mL) was used as the secondary antibody, resuspended in 0.5% nonfat milk in 1x TBS-T. 2 µL of LI-COR IRDye 680RD Goat anti-mouse IgG (1,000 µg/mL) was used as the secondary antibody of the loading control. Washes were done with 1x TBS-T, with the final wash being 1x TBS. Blots were analyzed at 700 and 800 nm wavelengths using the LI-COR Odyssey Fc Imager.

### C-di-GMP reporter activity quantification

Liquid LBS was inoculated with 5 μL of glycerol stock of each strain from an arrayed strain collection and grown at room temperature (23-25°C) overnight on a shaker. 2 or 4 μL of liquid culture were spotted onto LBS agar and incubated at 25°C for 24 h. For assay of strains not in 96-well format (**FIG. S2B and C**), strains were streaked on LBS agar and single colonies were inoculated into liquid LBS and grown at room temperature (23-25°C) overnight on a shaker. 4 or 8 μL of liquid culture were spotted onto LBS agar and incubated at 25°C for 24 h. For both versions of the assay, spots were resuspended in 500 μL 70% Instant Ocean (IO), then OD_600_, TurboRFP (555 nm excitation/585 nm emission), and AmCyan (453 nm excitation/486 nm emission) for each resuspended spot were measured in triplicate using BioTek Synergy Neo2 plate reader. To calculate c-di-GMP reporter activity, TurboRFP values (reports on c-di-GMP) were normalized to AmCyan values (constitutively expressed).

### ELISA c-di-GMP quantification

Overnight cultures were spotted onto LBS agar and incubated at 25°C overnight. Spots were resuspended in UltraPure water (Cayman Chemical), pelleted in a table top microcentrifuge, and frozen at −80°C. To standardize cell density, Bacteria Protein Extraction Reagent (B-PER, Thermo Fisher Scientific) was added to cell pellets at a ratio of 1:4 w/v then incubated for 10 min at room temperature (23-25°C). Cell debris was pelleted and the supernatants were diluted for c-di-GMP quantification. Samples were processed and quantified using the Cyclic di-GMP ELISA Kit by Cayman Chemical (Item #501780). Absorbance of the samples were measured at 450 nm using a BioTek Synergy Neo2 plate reader to determine c-di-GMP concentration. A standard curve was generated using the provided Cayman Chemical ELISA Analysis Tool, followed by sample quantification using the standard curve.

### Data analysis

Congo red binding was quantified using ImageJ (version 2.0.0-rc-69/1.52p) by subtracting the WT gray value from the mutant gray value and multiplying the value by −1 (36). Fluorescence of reporter strains in liquid culture and ELISA samples were measured using a BioTek Synergy Neo2 plate reader. Western blots were imaged using the LI-COR Odyssey Fc Imager. GraphPad Prism was used to generate graphs and conduct statistical analyses. Graphs were further refined in Adobe Illustrator.

## ACKNOWLEDGMENTS

We are grateful to Ella Rotman and John Brooks for early contributions to the project; Ketan Kotla for active site mutant construction; and Chris Waters for the pEVS143-QrgB and pEVS143-VC1086 plasmids. This study was funded by NIGMS grant R35 GM148385 to M.J.M., and R.Y.I. was supported by NIGMS training grant T32 GM007215.

---

**FIG S1. Many predicted *V. fischeri* DGCs and PDEs impact swimming motility when overexpressed. A.** Summary of motility results for overexpression of indicated proteins in TBS, TBS-Mg, and TBS-Ca soft (0.3%) agar. Blue coloring indicates phenotypes expected from elevated c-di-GMP, whereas pink indicates phenotypes expected from reduced c-di-GMP. White indicates no significant change. **B.** Quantification of migration through TBS, TBS-Mg, and TBS-Ca soft (0.3%) agar for *V. fischeri* strains overexpressing the indicated proteins relative to the pRYI039 empty vector control. TBS data is the same as represented in FIG 2B. For each strain, n = 4-12 biological replicates (33-36 for controls). One-way analysis of variance (ANOVA) was used for statistical analysis, each bar represents the means of biological replicates, error bars represent standard errors of the mean, asterisks represent significance relative to the pRYI039 empty vector control (*, *P* < 0.05), and numbers represent VF_ locus tags (e.g., VF_0087, VF_A0056, etc.); negative controls pRYI039 and pEVS143 as well as non-*V. fischeri* controls QrgB and VC1086 are also listed and indicated with a black dot.

**FIG S2. C-di-GMP biosensor activity.** Quantification of c-di-GMP concentration for *V. fischeri* strains overexpressing the indicated proteins using the pFY4535 c-di-GMP reporter plasmid. **A.** Assays performed in 96-well format. Values are relative to the pRYI039 empty vector control. For each strain, n = 3-6 (15 for control) biological replicates. **B.** Assays performed in non-96-well format. Values are relative to the pRYI039 empty vector control. For each strain, n = 3-7 biological replicates (11 for control). **C.** Assays of indicated proteins overexpressed in a high c-di-GMP background. For each strain, n = 3-9 biological replicates. **D.** Integration of phenotypic data for the *V. fischeri* DGCs and PDEs. Blue coloring indicates phenotypes expected from elevated c-di-GMP, whereas pink indicates phenotypes expected from reduced c-di-GMP. White indicates no significant change. For A to C, constitutive AmCyan was used to normalize TurboRFP to cell density, one-way analysis of variance (ANOVA) was used for statistical analysis, each bar represents the means of biological replicates, error bars represent standard errors of the mean, asterisks represent significance relative to the pRYI039 empty vector control (*, P < 0.02). For A to D, numbers represent VF_ locus tags (e.g., VF_0087, VF_A0056, etc.); negative controls pRYI039 and pEVS143 as well as non-*V. fischeri* controls QrgB and VC1086 are also listed and indicated with a black dot.

**FIG. S3. C-di-GMP quantification methods do not match PDE functional characterization. A.** Quantification of Congo red binding for *V. fischeri* PDEs overexpressing the indicated proteins relative to the pRYI039 empty vector control. For each strain, n = 5 biological replicates. Each bar represents the means of biological replicates. Data are the same as those represented in FIG 2A. **B.** Quantification of migration through soft (0.3%) agar for *V. fischeri* PDEs overexpressing the indicated proteins relative to the pRYI039 empty vector control. For each strain, n = 4 biological replicates. Each bar represents the means of biological replicates. Data are the same as those represented in FIG 2B. **C.** Quantification of c-di-GMP concentration for *V. fischeri* strains overexpressing the indicated PDEs using the pFY4535 c-di-GMP reporter plasmid (left y-axis; open dots) and ELISA (right y-axis; solid dots). Values are relative to the pRYI039 empty vector control. For each strain, n = 3 (9 for controls) biological replicates, dots represent the means of technical replicates, average bars represent the means of biological replicates. C-di-GMP reporter data are the same as those represented in FIG S2A. For A-C, error bars represent standard errors of the mean and numbers represent VF_ locus tags (e.g., VF_0087, VF_A0056, etc.); negative control pRYI039 and non-*V. fischeri* control VC1086 are also listed and indicated with a black dot.

