## Supplementary material for "Functional analysis of cyclic diguanylate-modulating proteins in *Vibrio fischeri*": Figures S1-S3

**A**

| | Protein | Motility TBS | Motility Mg | Motility Ca |
|---|---|---|---|---|
| DGCs (GGDEF) | 0596 (CdgG) | | | |
| | 0989 (MifA) | - | - | - |
| | 1200 | - | - | - |
| | 1245 | | | |
| | 1350 | - | | |
| | 1515 | | | |
| | 1561 | - | - | - |
| | 1639 (CasA) | - | - | - |
| | 2261 | - | - | |
| | 2362 | - | - | - |
| | A0056 | | | |
| | A0057 | - | - | - |
| | A0152 | - | - | - |
| | A0155 | - | - | - |
| | A0276 | - | - | - |
| | A0323 | - | - | - |
| | A0342 | - | - | - |
| | A0343 | - | - | - |
| | A0368 | - | - | - |
| | A0381 | - | - | - |
| | A0398 | - | - | - |
| | A0476 | | | |
| | A0567 | - | - | - |
| | A0692 | - | - | - |
| | A0796 | | | |
| | A0959 (MifB) | - | - | - |
| | A0976 | | | + |
| | A1012 | - | - | - |
| | QrgB | - | - | - |
| PDEs (EAL) | 0087 | | | |
| | 0091 | | | |
| | 1603 | + | | + |
| | 2480 | + | | + |
| | A0344 | | | + |
| | A0526 | | | |
| | A0551 | + | | |
| | A0706 | | | |
| | A0879 | | | |
| | A1014 (PdeV) | | | |
| | A1038 (BinA) | + | | + |
| | A1076 | | | |
| | VC1086 | + | | + |
| PDEs (HD-GYP) | 1367 | | | |
| | A0506 | + | | + |
| DGC/PDEs | 0094 | + | | + |
| | 0494 | + | | + |
| | 0985 | - | | - |
| | A0244 | | | |
| | A0475 | | | |
| Degenerate | 0355 | | | |
| | A0216 | | | |
| | A1166 (LapD) | - | - | |

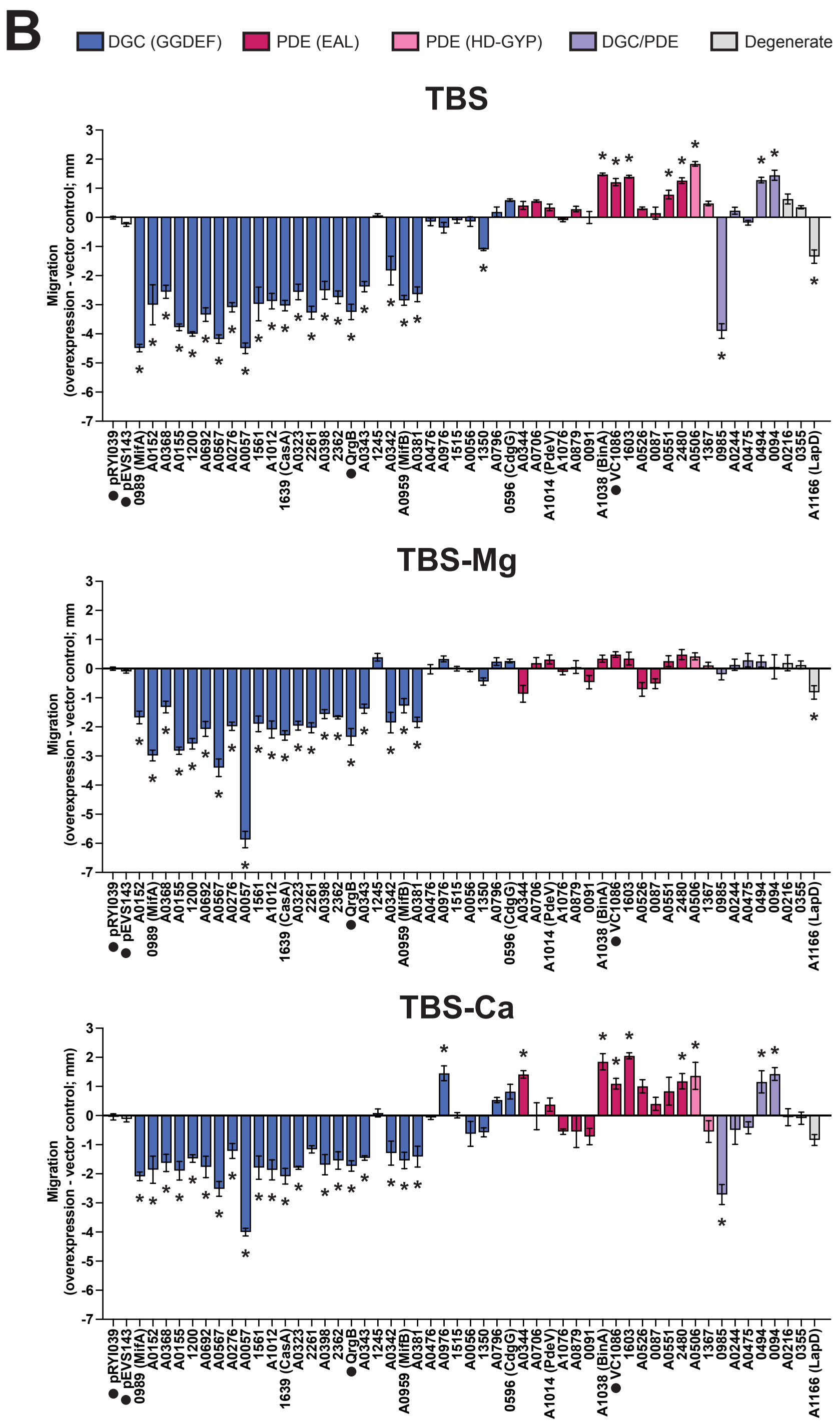


**FIG S1. Many predicted V. fischeri DGCs and PDEs impact swimming motility when overexpressed.** **A.** Summary of motility results for overexpression of indicated proteins in TBS, TBS-Mg, and TBS-Ca soft (0.3%) agar. Olive coloring indicates phenotypes expected from elevated c-di-GMP, whereas orange indicates phenotypes expected from reduced c-di-GMP. White indicates no significant change. **B.** Quantification of migration through TBS, TBS-Mg, and TBS-Ca soft (0.3%) agar for *V. fischeri* strains overexpressing the indicated proteins relative to the pRYI039 empty vector control. TBS data is the same as represented in FIG 2B. For each strain, n = 4-12 biological replicates (33-36 for controls). One-way analysis of variance (ANOVA) was used for statistical analysis, each bar represents the means of biological replicates, error bars represent standard errors of the mean, asterisks represent significance relative to the pRYI039 empty vector control (*, $P < 0.05$), and numbers represent VF_ locus tags (e.g., VF_0087, VF_A0056, etc.); negative controls pRYI039 and pEVS143 as well as non-*V. fischeri* controls QrgB and VC1086 are also listed and indicated with a black dot

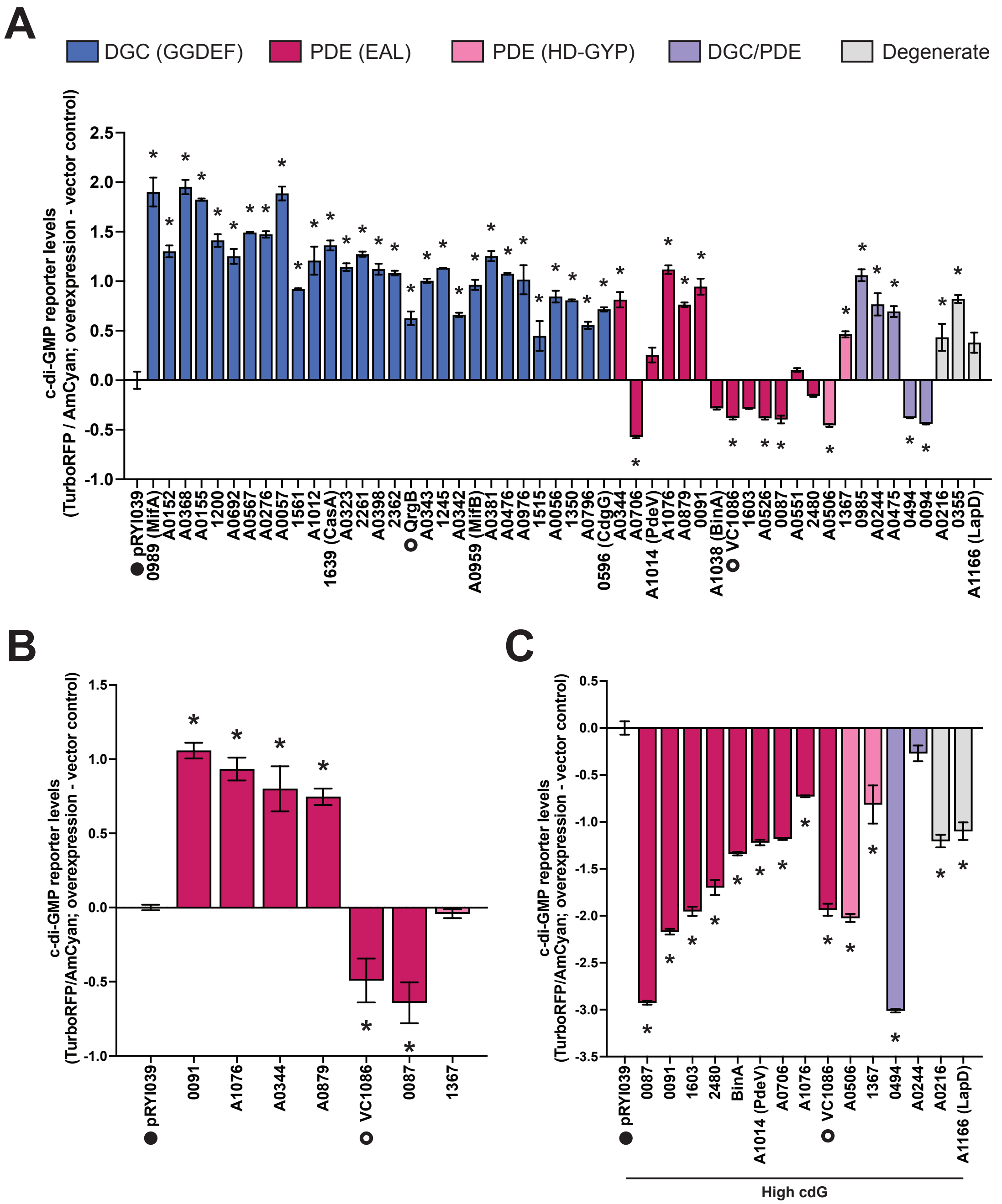


**D**

| Protein | cdG | CR | Mot |
|---|---|---|---|
| 0596 | + | | |
| 0989 (MifA) | + | + | - |
| 1200 | + | + | - |
| 1245 | + | | |
| 1350 | + | | - |
| 1515 | + | | |
| 1561 | + | + | - |
| 1639 (CasA) | + | + | - |
| 2261 | + | + | - |
| 2362 | + | + | - |
| A0056 | + | | |
| A0057 | + | + | - |
| A0152 | + | + | - |
| A0155 | + | + | - |
| A0276 | + | + | - |
| A0323 | + | + | - |
| A0342 | + | | - |
| A0343 | + | + | - |
| A0368 | + | + | - |
| A0381 | + | | - |
| A0398 | + | + | - |
| A0476 | + | | |
| A0567 | + | + | - |
| A0692 | + | + | - |
| A0796 | + | | |
| A0959 (MifB) | + | | - |
| A0976 | + | | |
| A1012 | + | + | - |
| QrgB | + | + | - |
| 0087 | - | - | |
| 0091 | + | - | |
| 1603 | | - | + |
| 2480 | | - | + |
| A0344 | + | | |
| A0526 | - | - | |
| A0551 | | - | + |
| A0706 | - | | |
| A0879 | + | | |
| A1014 (PdeV) | | | |
| A1038 (BinA) | | - | + |
| A1076 | + | | |
| VC1086 | - | - | + |
| 1367 | + | - | |
| A0506 | - | - | + |
| 0094 | - | - | + |
| 0494 | | - | + |
| 0985 | + | + | - |
| A0244 | + | | |
| A0475 | + | | |
| 0355 | + | | |
| A0216 | + | | |
| A1166 (LapD) | | | - |

**FIG S2. C-di-GMP biosensor activity.** Quantification of c-di-GMP concentration for *V. fischeri* strains overexpressing the indicated proteins using the pFY4535 c-di-GMP reporter plasmid. **A.** Assays performed in 96-well format. Values are relative to the pRYI039 empty vector control. For each strain, n = 3-6 (15 for control) biological replicates. **B.** Assays performed in non-96-well format. Values are relative to the pRYI039 empty vector control. For each strain, n = 3-7 biological replicates (11 for control). **C.** Assays of indicated proteins overexpressed in a high c-di-GMP background. For each strain, n = 3-9 biological replicates. **D.** Integration of phenotypic data for the *V. fischeri* DGCs and PDEs. Olive coloring indicates phenotypes expected from elevated c-di-GMP, whereas red-orange indicates phenotypes expected from reduced c-di-GMP. White indicates no significant change. For A to C, constitutive AmCyan was used to normalize TurboRFP to cell density, one-way analysis of variance (ANOVA) was used for statistical analysis, each bar represents the means of biological replicates, error bars represent standard errors of the mean, asterisks represent significance relative to the pRYI039 empty vector control (*, P < 0.02). For A to D, numbers represent VF_ locus tags (e.g., VF_0087, VF_A0056, etc.); non-*V. fischeri* controls QrgB and VC1086 are also listed and indicated with an open circle while negative controls are indicated with a closed circle.

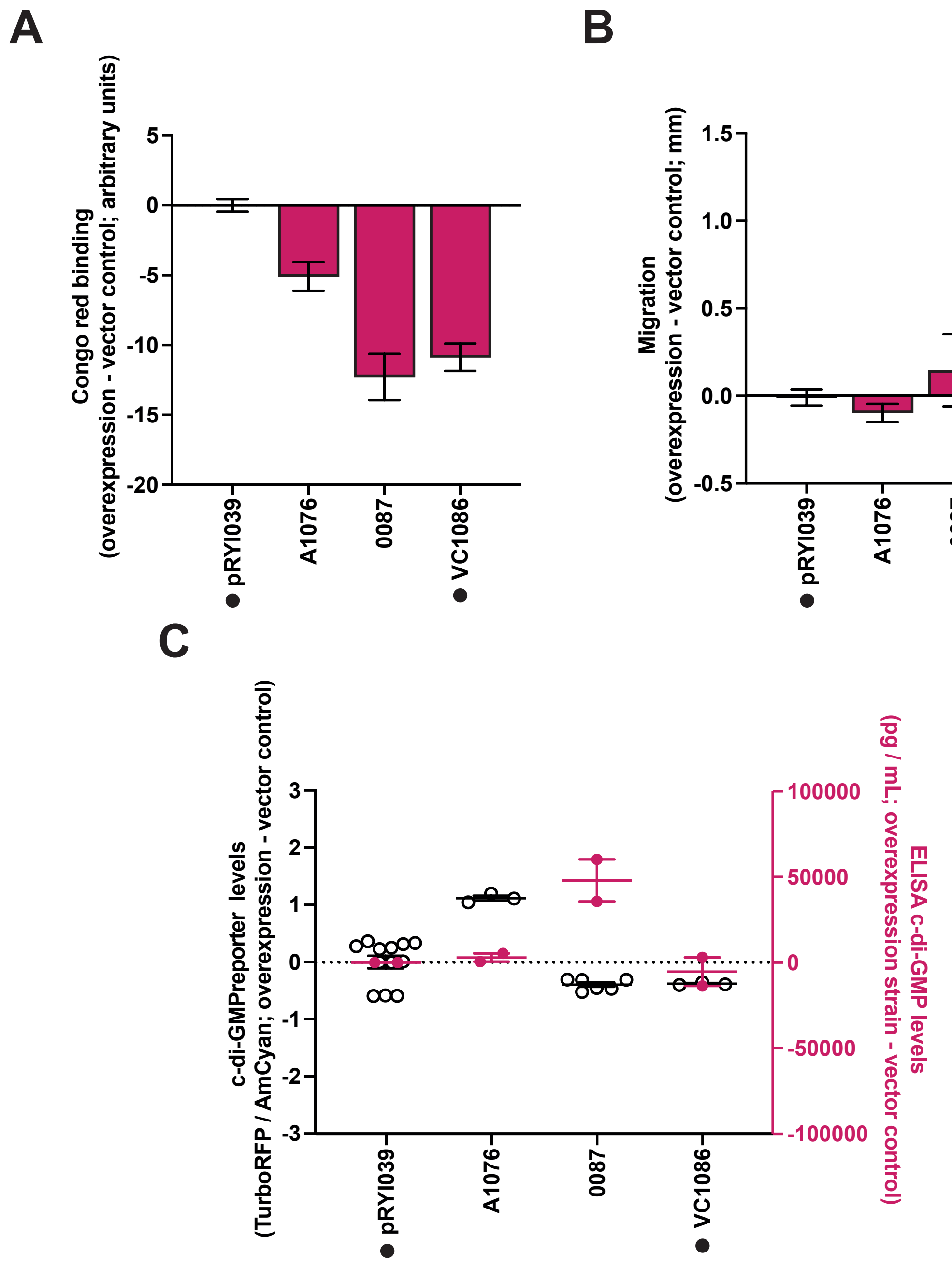


**FIG. S3. C-di-GMP quantification methods do not match PDE functional characterization.**
**A.** Quantification of Congo red binding for *V. fischeri* PDEs overexpressing the indicated proteins relative to the pRYI039 empty vector control. For each strain, n = 5 biological replicates. Each bar represents the means of biological replicates. Data are the same as those represented in FIG 2A. **B.** Quantification of migration through soft (0.3%) agar for *V. fischeri* PDEs overexpressing the indicated proteins relative to the pRYI039 empty vector control. For each strain, n = 4 biological replicates. Each bar represents the means of biological replicates. Data are the same as those represented in FIG 2B. **C.** Quantification of c-di-GMP concentration for *V. fischeri* strains overexpressing the indicated PDEs using the pFY4535 c-di-GMP reporter plasmid (left y-axis; open dots) and ELISA (right y-axis; solid dots). Values are relative to the pRYI039 empty vector control. For each strain, n = 3 (9 for controls) biological replicates, dots represent the means of technical replicates, average bars represent the means of biological replicates. C-di-GMP reporter data are the same as those represented in FIG S2A. For A-C, error bars represent standard errors of the mean and numbers represent VF_ locus tags (e.g., VF_0087, VF_A0056, etc.); negative control pRYI039 and non-*V. fischeri* control VC1086 are also listed and indicated with a black dot.
